# TNFa and IL-6 promote ex-vivo proliferation of lineage-committed human regulatory T cells

**DOI:** 10.1101/2021.08.09.455690

**Authors:** Nikolaos Skartsis, Yani Peng, Leonardo M.R. Ferreira, Vinh Nguyen, Yannick Muller, Flavio Vincenti, Qizhi Tang

## Abstract

Treg therapy is being tested in clinical trials in transplantation and autoimmune diseases, however, the impact of inflammation on Tregs is unclear. In this study, we challenged human Tregs ex-vivo with pro-inflammatory cytokines, TNFα and IL-6. These cytokines enhanced Treg proliferation induced by anti-CD3 and anti-CD28 or CD28 superagonist (CD28SA) while maintaining high expression of FOXP3 and HELIOS, demethylated FOXP3 enhancer, and low expression of cytokines IFNg, IL-4 and IL-17. Blocking TNF receptor signaling using etanercept or deletion of TNF receptor 2 using CRISPR/Cas9 blunted Treg proliferation and attenuated FOXP3 and HELIOS expression, revealing the importance of TNFR2 signaling in Treg proliferation and lineage stability. The robust proliferation induced by CD28SA with IL-6 and TNFα may be adopted for the expansion of therapeutic Tregs. Metabolomics analysis showed that Tregs expanded with CD28SA plus cytokines had more active glycolysis and oxidative phosphorylation, increased energy production, and higher antioxidant potential. Finally, CD28SA plus cytokine-expanded Tregs had comparable suppressive activity in vitro and in vivo in a humanized mouse model of graft-versus-host-disease when compared to Tregs expanded using the conventional protocol. These results demonstrate that human Tregs positively respond to proinflammatory cytokines with enhanced proliferation without compromising their lineage identity or function.

## Introduction

Tregs are a subset of CD4^+^ T cells dedicated to maintaining normal immune homeostasis (1). These cells can be identified by the expression of the transcription factor Forkhead box protein P3 (FOXP3), a master regulator of Treg lineage specification and Treg function (2). Tregs prevent unwanted immune activation in steady state and promote resolution of immune response at the site of inflammation (3-5). Tregs are frequently enriched among inflammatory infiltrates. Their function in suppressing immune system activation and promoting tissue repair is supported by numerous experimental and clinical observations (6-8). In mouse models of type 1 diabetes, Tregs isolated from inflamed islets, but not those from lymphoid organs, are able to suppress anti-islet autoimmune attacks (9). Similarly, in a mouse model of multiple sclerosis, Tregs in the central nervous system (CNS) are critical for limiting pathology at the peak of disease and this tissue-restricted function of Tregs depends on TNF receptor 2 (TNFR2) signaling (10). In a mouse model of islet transplant, it was reported that Tregs first traffic to the inflamed graft to suppress alloimmune responses and subsequently retreat to draining lymph nodes to maintain tolerance (11). In humans, higher FOXP3 in graft biopsies predicts reversibility of T cell-mediated rejection of kidney transplant whereas higher Tregs in the tumor microenvironment is associated with worse survival in most cancers (12, 13). However, it has been suggested that Tregs may be destabilized under certain inflammatory conditions and there has been considerable debate on the impact of proinflammatory signals on Treg function (14). During immune activation, proinflammatory cytokines such as TNFα and IL-6 are produced by various cells including macrophages, neutrophils, T cells, B cells, and stromal cells (15-18). TNFα and IL-6 promote T cell activation (19, 20) and increase resistance to Treg-mediated suppression (21-23), both of these functions contribute to mounting an effective immune response. While there is consensus that TNFα and IL-6 heighten effector T cell (Teff) resistance to suppression by Tregs (24, 25), their direct impact on Tregs, especially on human Tregs, are less clear.

The effect of TNFα on mouse Tregs have been extensively studied. While TNFα impairs the differentiation and function of TGFβ-induced Tregs (26), it is generally accepted that TNFα exerts positive effects on thymic derived Tregs (27-29). TNFα optimally activates Tregs via upregulating TNFR2, 4-1BB, and OX40 expression, which leads to increased Treg proliferation (30, 31). TNFR2-deficient Tregs lose Foxp3 expression and are unable to protect against colitis when co-transferred with naïve CD4^+^ T cells into Rag1^KO^ mice (32). Conditional inactivation of TNFR2 in Tregs leads to reduced expression of Foxp3, CD25, cytotoxic T-lymphocyte-associated protein 4 and Glucocorticoid-induced tumor necrosis factor receptor and exacerbates experimental autoimmune encephalitis (EAE) (33). TNFR2-expressing Tregs are critical to suppressing EAE by limiting T cell activation in the CNS during overt inflammation (10). In addition, TNFR2 expression defines a maximally suppressive subset of Tregs that accumulates intratumorally in a Lewis lung carcinoma model (34). Finally, TNFα-primed mouse Tregs have enhanced efficacy against graft-versus-host disease (GVHD) when compared with unprimed Tregs (35, 36). Overall, these results suggest that TNFα negatively affects TGFβ-induced Tregs, but exerts a positive effect on thymic derived Treg activation and function in mice.

The effects of TNFα on human Tregs has been more controversial. Tregs isolated from rheumatoid arthritis patients have been reported to lose FOXP3 expression and convert into pathogenic T cells when exposed to TNFα in vitro (37, 38). TNFα partially abrogated the suppressive function of CD4^+^CD25^+^ T cells isolated from chronic HBV-infected patients (39). Activation of the canonical NF-κB pathway in CD45RA^-^ Tregs by TNFα-TNFR2 interaction downmodulated their suppressive function (40). However, these findings have been challenged by others who report a pro-regulatory role of TNFα in optimally activating Tregs isolated from healthy donors, without negatively affecting their in vitro suppressive function (41). Similar to studies in mouse Tregs, TNFR2 expression identifies a subset of highly suppressive Tregs isolated from PBMC (42). TNFR2 signaling has been shown to enhance human Treg proliferation in response to IL-2 via activation of the non-canonical NF-κB pathway (43). As a result, TNFR2 agonists are used in ex-vivo human Treg expansion protocols to facilitate homogeneous high purity cellular products (44, 45). Multiple studies have shown that TNFα blockade therapy paradoxically exacerbate autoimmune pathologies in psoriasis and demyelinating diseases (46, 47), suggesting a regulatory role of TNFα in these disease settings.

Multiple studies reported a negative role of IL-6 on mouse Tregs. IL-6 prevents the differentiation of naïve CD4^+^ T cells into the peripherally induced Tregs in a dominant and non-redundant manner (48). IL-6 has been reported to destabilize mouse thymic derived Tregs and drive them to produce IL-17 in vitro (49). However, Tregs isolated from inflamed CNS lesions in EAE-affected mice are insensitive to IL-6-driven Th17 conversion mainly due to a downregulation of CD126 (IL-6Ra) and CD130 (gp130), the signaling component of the IL-6R complex (50). IL-6 has been found to increase the rate of Foxp3 proteasomal degradation via downregulation of the expression of deubiquitinase USP7, which is normally upregulated and associated with Foxp3 in the nucleus of Tregs (51). In addition, IL-6-induced downregulation of Foxp3 expression is attenuated in Deleted in breast cancer 1-deficient Tregs due to decreased Caspase 8-mediated Foxp3 degradation (52). In contrast to the aforementioned studies, IL-6 Tg mice harbor increased numbers of Tregs in their lymphoid organs with intact functional capacity against naïve T cells in vitro (53). While IL-6 enhances the generation of Th17 cells, classic signaling via IL-6Ra induces double positive RORγt^+^Foxp3^+^ Tregs with potent immunoregulatory properties in a mouse model of glomerulonephritis (54). The current prevailing view is that IL-6 negatively impacts mouse Tregs, but emerging data suggest that differential mode of signaling (classic versus trans-signaling) may have divergent impacts on mouse Tregs.

Similar to findings in mouse Tregs, IL-6 prevents in-vitro induction of human Tregs from conventional T cells (Tconvs) (55). In addition, peripheral blood CD4^+^CD25^+^ Tregs downregulate FOXP3 and lose their suppressive function following IL-6 exposure (56). Retinoic acid, known for its stabilizing effects on induced Tregs and thymic Tregs, downregulates IL-6Ra and renders Tregs insensitive to the destabilizing effect of IL-6 (56). gp130^hi^ Tregs are enriched amongst naïve CD45RA^+^ Tregs isolated from human PBMC and have lower in vitro suppressive capacity when compared to gp130^lo^ Tregs (57). These results support the notion that IL-6 negatively impacts human Tregs. Another group has immunophenotyped CD4^+^CD127^lo^CD25^+^ Tregs isolated from PBMC and reported the presence of a subpopulation of human IL-6Ra^hi^CD45RA^-^ Tregs that have potent suppressive capacity in vitro, demethylated Treg-specific demethylated region (TSDR), a Th_17_ transcriptional signature, and produce both pro- and anti-inflammatory cytokines, such as IL-17, IL-22, and IL-10 (58). These studies highlight the heterogeneity of circulating human Tregs. Overall, there is a relative paucity of studies measuring the direct impact of IL-6 on human Treg proliferation, phenotypic stability, and function.

In this study, we examined the direct impact of TNFα and IL-6 on human Tregs in an ex-vivo culture system and evaluated in-vivo function of TNFα and IL-6 exposed human Tregs in a xenogeneic GVHD model in humanized mice. Our data show a positive role of both IL-6 and TNFα on Tregs in promoting their proliferation without lineage destabilization, suggesting that Tregs can respond to proinflammatory signals to increase their presence at sites of inflammation without compromising their lineage stability.

## Results

### TNFα and IL-6 promoted proliferation of lineage-committed Tregs

To determine the direct effects of TNFα and IL-6 on human Tregs, we challenged FACS purified CD4^+^CD25^+^CD127^lo/-^ Tregs with IL-6 and/or TNFα, during in vitro activation. We stimulated the cells with anti-CD3 and anti-CD28 (aCD3/28) beads in the presence of 300 IU/ml recombinant human IL-2 to simulate antigen activation of Tregs. Activated Tregs formed multicellular bead and cell clusters, which is followed by cell proliferation and return to a single-celled quiescent state by day 9 of culture as previously reported (59). The addition of IL-6, TNFα, or both during the first 5 days of the Treg culture resulted in more persistent clustering of the cells (Figure 1A). Moreover, combination of IL-6 and TNFα resulted in significantly improved Treg expansion, which mainly manifested during the later stage of the culture (Figure 1B).

**Figure 1.**
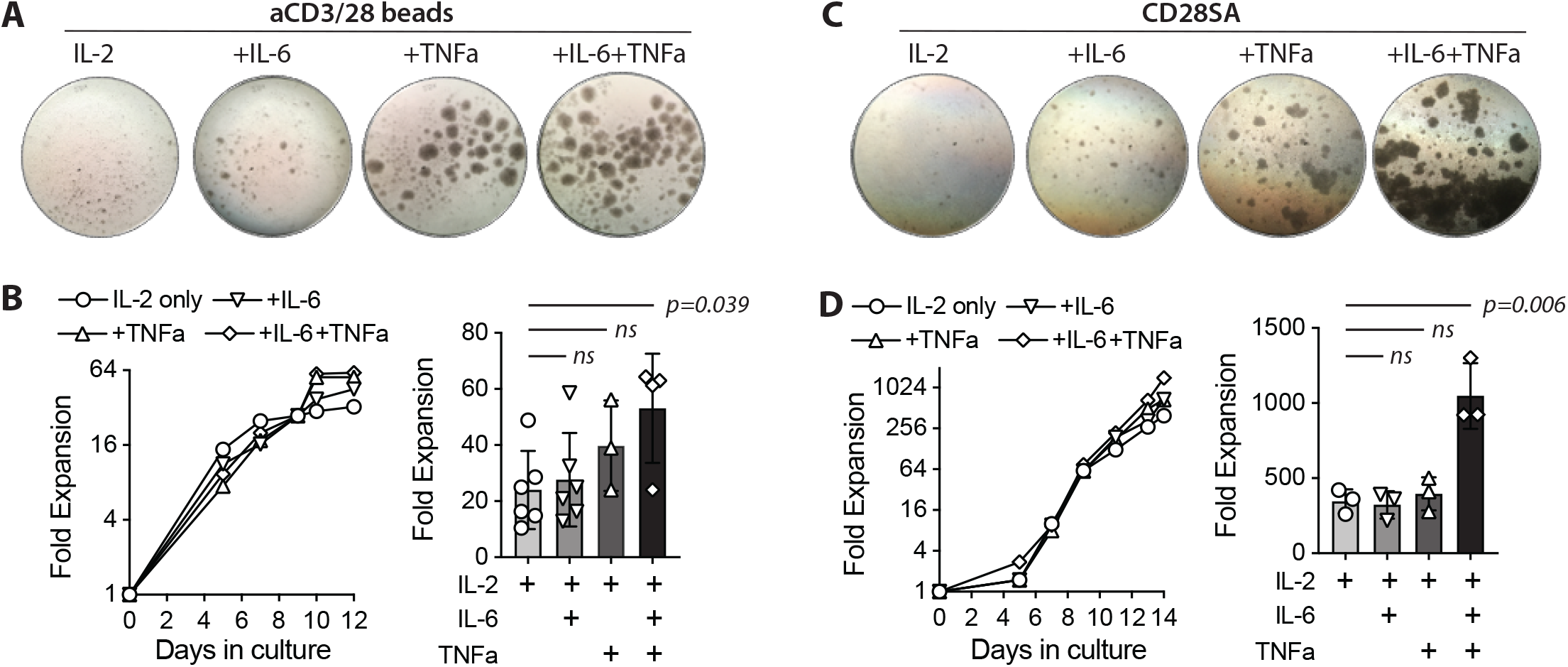
TNFa and IL-6 augmented ex-vivo proliferation of human Tregs. FACS purified human Tregs were stimulated with either aCD3/28 beads or CD28SA and cultured in the presence of 300 IU/ ml rhIL-2 with or without TNFa and IL-6 for 9 and 14 days as indicated (**A**) Microscopic pictures of aCD3/28 bead-stimulated Treg cultures on day 8 of culture. Original magnification was 10x. (**B**) An example of aCD3/28 bead-stimulated Treg expansion kinetics (left) and a summary of fold expansion on day 9 of 3 to 6 independent experiments (right) are shown. (**C** and **D**) Same as panel A and B, except the Tregs were stimulated with CD28SA. Statistical significance of the differences was assessed using one-way ANOVA and Dunnett’s multiple comparisons posttest using IL-2 alone as baseline references. p values are stated. Results shown used cells from 3 distinct donors and are representative of 6 independent experiments using cells from 6 distinct donors.

It has been previously reported that CD28 superagonist antibodies (CD28SA) can also induce ex-vivo expansion of human Tregs (60). We thus replaced the aCD3/28 beads with CD28SA and evaluated the impact of IL-6 and TNFα in the context of a different mitogenic stimulation. While CD28SA-stimulated Tregs showed delayed kinetics in proliferation when compared to their aCD3/CD28-stimulated counterparts, the cells proliferated more persistently after a single stimulation, resulting in more overall expansion (Figure 1C and 1D). Exposure to both TNFα and IL-6 resulted in robust cell clustering over a prolonged period and synergistic enhancement of cell expansion that was far more pronounced than seen with aCD3/CD28 bead-stimulated Tregs (Figure 1B and 1D).

Since FOXP3 had been reported to limit Treg proliferation and metabolism (61), we wondered if the enhanced proliferation in the presence of TNFα and IL-6 was due to the destabilization of Tregs into FOXP3-negative exTregs. We measured the expression of FOXP3 and HELIOS, the two lineage-defining transcription factors for Tregs (62), at the end of the 9-day cell expansion. The results showed that Tregs exposed to IL-6 and/or TNFα maintained high levels of FOXP3 and HELIOS expression when compared to expanded Tconvs (Figure 2A and 2B). Activated human Tconvs can increase FOXP3 expression transiently, but their genomic DNA is methylated at the TSDR, which is demethylated in Tregs. Thus, demethylated TSDR is a more discriminating marker for bona fide Tregs. Therefore, we analyzed TSDR methylation status in Tregs cultured in various conditions. The results showed a high degree of TSDR demethylation in all expanded Tregs (Figure 2C), supporting the flow cytometric data to show that the exposure to IL-6 and/or TNFα did not alter Treg identity.

**Figure 2.**
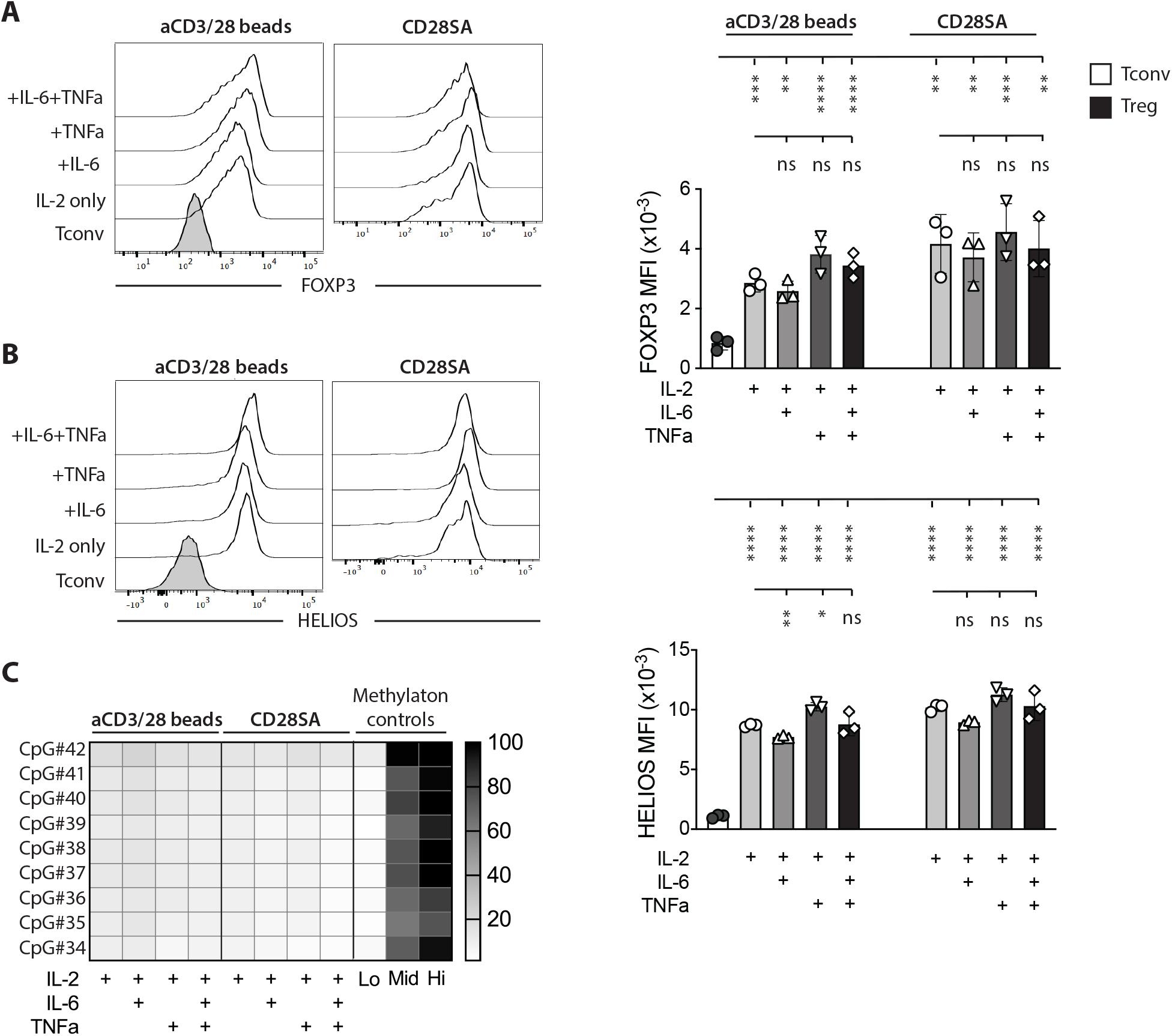
TNFa and IL-6 expanded Tregs maintained their lineage identity. FACS purified human Tregs were stimulated with either aCD3/28 beads or CD28SA and cultured in the presence of 300 IU/ml rhIL-2 with or without TNFa and IL-6 as indicated. (**A** and **B**) Flow cytometric analysis of FOXP3 (**A**) and HELIOS (**B**) expression in Tregs on day 9 after stimulation. Representative histograms (left) and summaries of MFI from 3 independent experiments (right) are shown. Statistical significance of differences was assessed using one-way ANOVA with Geisser-Greenhouse’s correction and Dunnett’s multiple comparisons test using either Tconv or Treg treated with IL-2 alone as a baseline reference. p values are marked as ns=not significant, p>0.05, *=p<0.05, **=p<0.01, ***=p<0.001, ****=p<0.0001. (**C**) Heatmap summary of TSDR demethylation of Tregs expand-ed in various conditions. Results shown are averages of Treg cultures using 2 unrelated male donors in 2 independent experiments.

To further assess if IL-6 and/or TNFα induced effector functions in Tregs, we measured the supernatant of Treg cultures on day 7 after stimulation for the presence effector cytokines using a 42-plex Luminex assay. Among the 42 cytokines in the panel, 21 were not present in any of the T cell cultures, 3 were added (IL-2, IL-6, and TNFα), and 18 were detected in Treg cultures. Among the 18 *de novo* cytokines detected in the culture, the amounts found in Treg cultures were markedly lower than those seen in the supernatant of Tconv cultures (Figure 3A).

**Figure 3.**
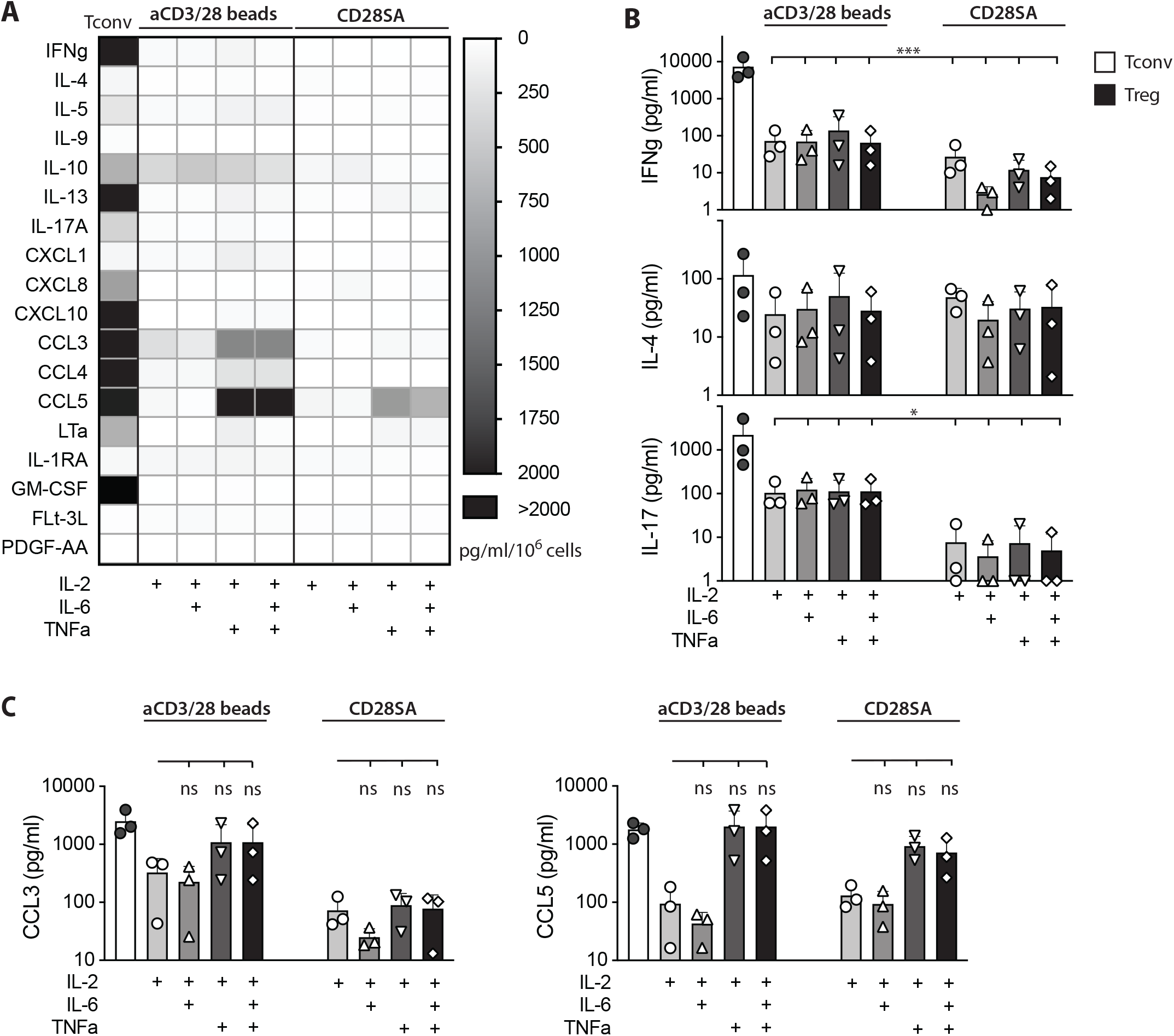
Tregs expanded in IL-6 and TNFa did not secrete more proinflammatory cytokines. FACS purified human Tregs were stimulated with either aCD3/28 beads or CD28SA and cultured in the presence of 300 IU/ml rhIL-2 with or without TNFa and IL-6 as indicated. Cytokine and chemokine secretion in the culture supernatant of various Treg cultures was assessed using a multiplex Luminex panel. Supernatant in aCD3/28 bead stimulated Tconv cultures are included as a reference. (**A**) Heatmap summary of cytokines and chemokines that were present in any of the culture condition is shown. (**B**) IFN-g, IL-4, and IL-17 concentrations in the Day 7 culture supernatants are shown. (**C**) CCL3 and CCL5 concentrations in the Day 7 culture supernatant are shown. Results shown are summaries of 3 independent experiments using cells from 3 unrelated donors. Statistical significance of differences was assessed using one-way ANOVA with Geisser-Greenhouse’s correction and Dunnett’s multiple comparisons posttest using Treg treated with IL-2 alone (Panels B and C) as a baseline reference. p values are marked as ns=not significant, p>0.05, *=p<0.05, **=p<0.01, ***=p<0.001, ****=p<0.0001.

Particularly, no increase in IFNγ (Th_1_), IL-4 (Th_2_), or IL-17A (Th_17_) were seen in aCD3/28 bead or CD28SA stimulated cultures by the exposures to IL-6 and/or TNFα (Figure 3B). In fact, IL-6 did not induce increased secretion of any of the cytokines. On the other hand, exposure to TNFα, alone or in the presence of IL-6, led to a consistent trend of increase of CCL3 and CCL5 across all biological replicates (Figure 3C). Taken together, these experiments revealed that IL-6 and TNFα exposure resulted in a significant enhancement of human Treg proliferation without inducing Treg lineage destabilization.

These results contradicted some of the previous reports of destabilizing effect of proinflammatory cytokines on Tregs (37, 38, 63, 64). Many of these previous studies stimulated Tregs in the presence of lower concentration of IL-2 than we used. IL-2 is an essential cytokine for Treg lineage commitment and maintenance. We hypothesized that the high concentration of IL-2 in our culture condition may have protected Tregs from the destabilizing effect of TNFα and IL-6. Therefore, we challenged the Tregs by reducing IL-2 concentration from 300 IU/ml to 15 IU/ml, which resulted in diminished cell clustering and reduced proliferation (data not shown). FOXP3 expression was reduced when compared to cells cultured in 300 IU/ml of IL-2 but remained significantly elevated when compared to that expressed by similarly expanded Tconvs (Supplemental Figure 1A). HELIOS expression was comparably high in Tregs cultured in 15 IU/ml and 300 IU/ml of IL-2 (Supplemental Figure 1B). More importantly, Tregs exposed to TNFα and IL-6 in the presence of low IL-2 maintained demethylated TSDR (Supplemental Figure 1C) and did not produce more effector cytokines (Supplemental Figure 2A), including IFN-γ, IL-4, or IL-17 (Supplemental Figure 2B) when compared to Tregs stimulated without these cytokines. Similar to cultures stimulated in the presence of high IL-2, co-culture with TNFα stimulated higher secretion of CCL5 in Tregs (Supplemental Figure 2C). Thus, IL-6 and TNFα did not induce Treg lineage destabilization even when IL-2 was limited.

### TNFR2 signaling promoted ex-vivo proliferation of human Tregs and contribute to preserving Treg identity

While analyzing cytokine secretion in Treg cultures, we noticed that TNFα was consistently detected in all cultures in higher amounts in aCD3/28 bead-stimulated cultures than in CD28SA-stimulated cultures (Figure 4A). The amount of TNFα in the supernatant was not affected by the concentration of IL-2 or the addition of IL-6. Moreover, another TNF superfamily (TNFSF) member, lymphotoxin a (LTα), was also consistently detected (Figure 4B). The presence of TNFα, but not IL-6, significantly increased the level of LTα in the aCD3/28 bead-stimulated Treg culture supernatant and a moderate trend of increased LTα in CD28SA-stimulated cultures (Figure 4B). Both of TNFα and LTα bind to TNFR1 and TNFR2 (65). It has been well-established that Tregs preferentially express TNFR2 (31), we thus investigated the TNFR2 expression of in vitro stimulated human Tregs. In vitro stimulation with either aCD3/28 beads or CD28SA, without the addition of IL-6 or TNFα, uniformly increased TNFR2 expression by day 8 of culture, although CD28SA induced upregulation was delayed when compared to that induced by aCD3/CD28 beads (Figure 4C). Thus, activated Tregs produced TNFα and LTα and also have increased expression of TNFR2, suggesting that TNFα and LTα may function as autocrine or paracrine factors for activated Tregs.

**Figure 4.**
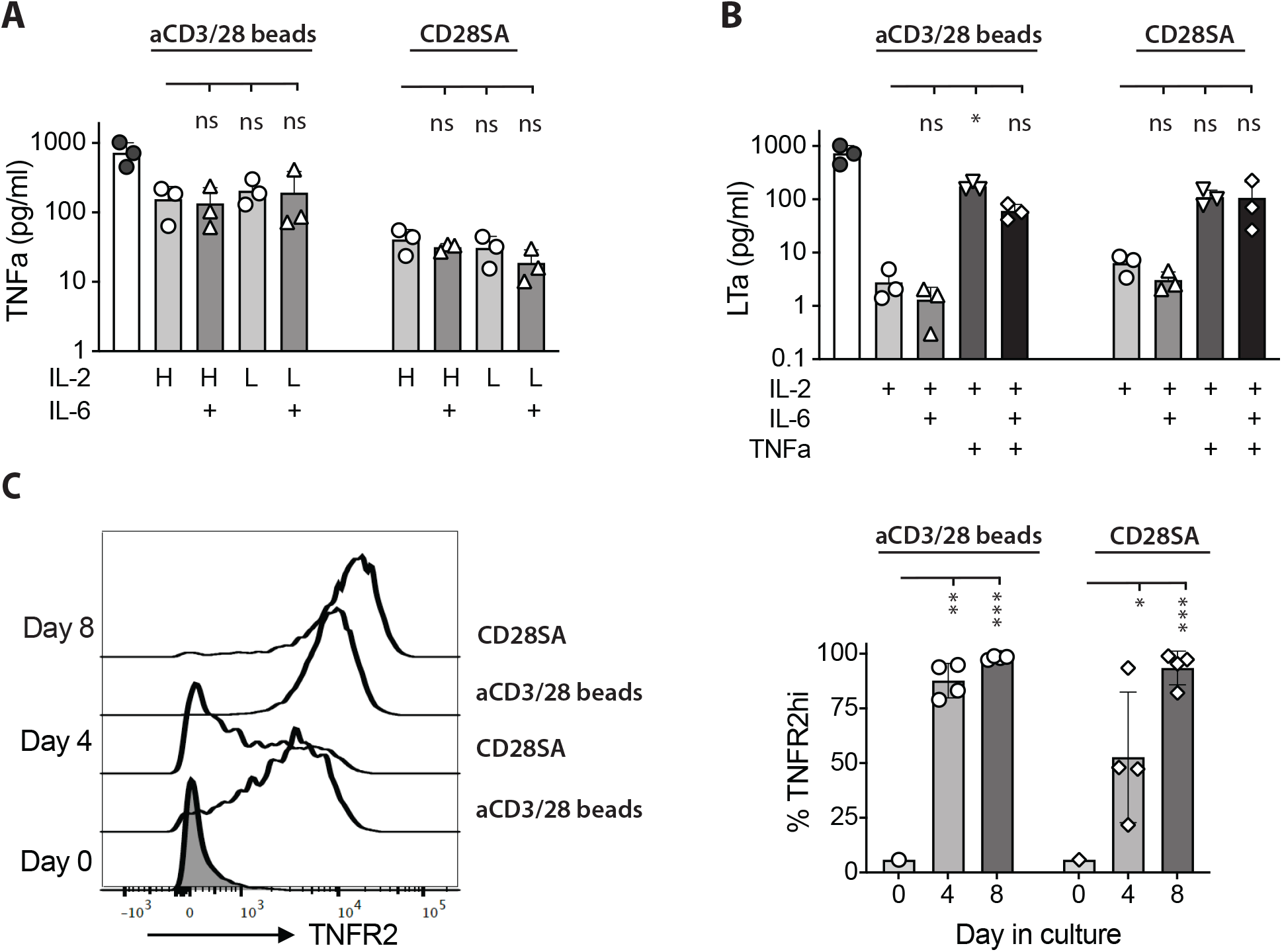
Activated Tregs produced TNFa and LTa and increased TNFR2 expression. FACS purified human Tregs were stimulated with either aCD3/28 beads or CD28SA and cultured in the presence of 300 IU/ml or 15 IU/ml rhIL-2 with or without IL-6 as indicated.(**A**) TNFa and (**B**) LTa concentration in the Day 7 culture supernatant of various T cell cultures. (**C**) Flow cytometric analysis of TNFR2 expression in Tregs stimulated with either aCD3/28 beads or CD28SA on day 0, 4 and 8 after stimulation. Representative histograms (left) and summary of percentage of TNFR2hi Tregs from 3 independent experiments (right) are shown. Statistical significance of differences was assessed using one-way ANOVA and Dunnett’s multiple comparisons posttest using Treg treated with IL-2 alone (Panels A and B) or Day 0 (Panel C) as a baseline reference. p values are marked as *=p<0.05, **=p<0.01, ***=p<0.001, ****=p<0.0001.

To investigate a potential role of TNFR2 in human Treg activation, we evaluated the impact of etanercept, a soluble TNFR2-Fc fusion protein, in ex-vivo Treg proliferation. Addition of etanercept throughout the duration of the culture significantly reduced Treg proliferation induced by either aCD3/CD28 beads or CD28SA (Figure 5A). The inhibitory effect of etanercept was most pronounced during the later stage of Treg culture, suggesting that TNFR2 signaling did not affect initial Treg proliferation likely due to delayed induction of TNFR2 or TNFα and LTα. Furthermore, etanercept significantly decreased expression of TNFR2 and the expanded Tregs showed a trend of moderate reduction in CD25, FOXP3, and HELIOS (Supplemental Figure 3).

**Figure 5.**
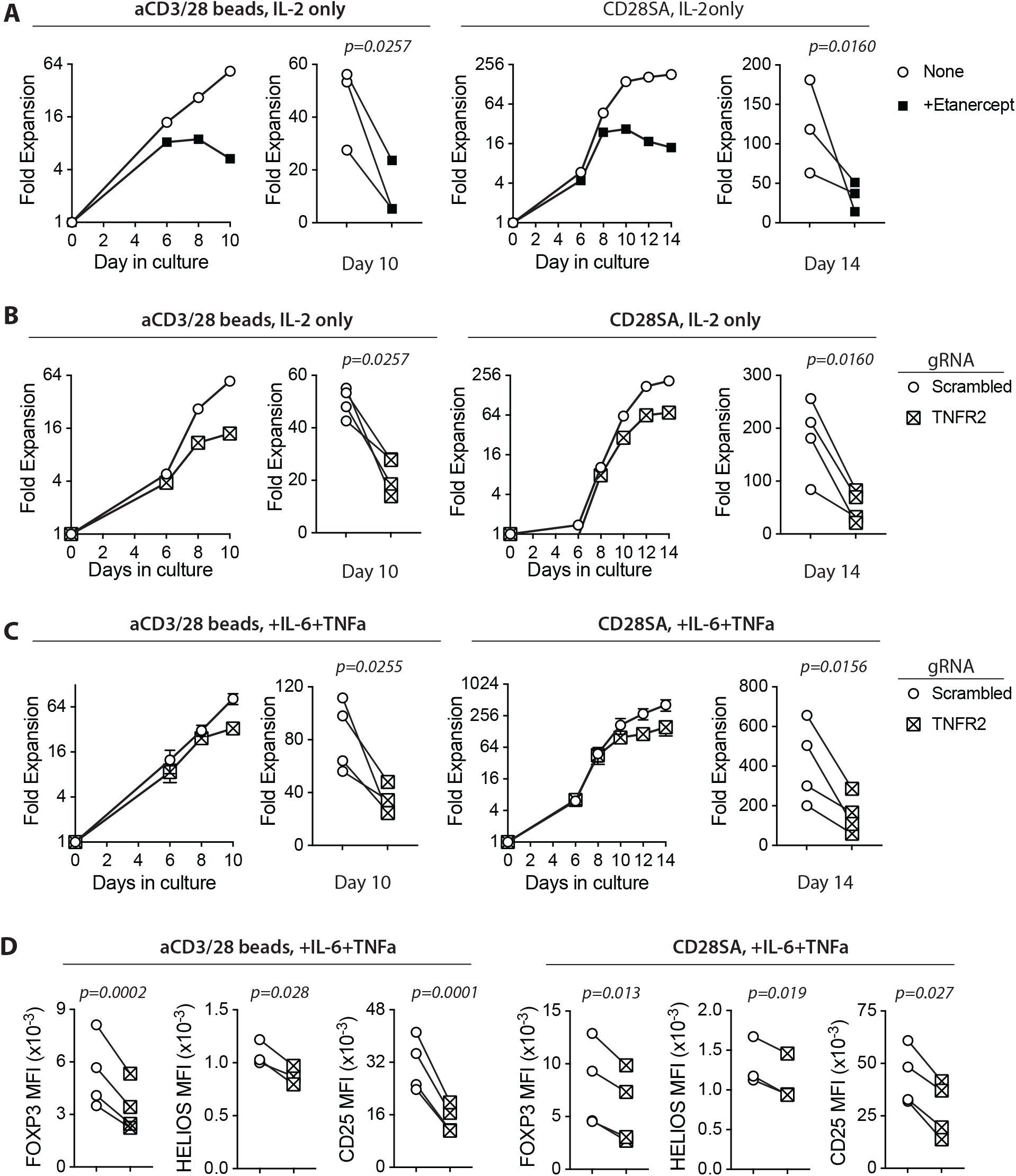
Ex-vivo proliferation of human Tregs is dependent on TNFR2 expression. FACS purified human Tregs were stimulated with either aCD3/28 beads or CD28SA and cultured in the presence of 300 IU/ml rhIL-2 with or without TNFa and IL-6 for 10 and 14 days, as indicated. (**A**) Representative aCD3/28 bead stimulated Treg expansion kinetics and summary of final fold expansion on day 10 of Treg expanded in the presence or absence of etanercept (5mcg/mL) (left). Similarly, CD28SA stimulated Tregs were expanded for 14 days (right). Results of 3 independent experiments using 3 unrelated donors are shown. (**B**) Tregs were gene edited to delete TNFR2 gene using CRISPR-Cas9 and then stimulated with either aCD3/28 beads (left) or CD28SA (right). Results shown are from 4 independent experiments using 4 unrelated donors. (**C**) Same as panel B, except the Tregs were cultured in the presence of IL-6 and TNFa. Results shown are from 4 independent donors in 4 independent experiments. (**D**) Tregs were expanded as shown in panel C and flow cytometric analysis of FOXP3, HELIOS and CD25 expression was performed on day 8 after stimulation. Results show summary of MFI of FOXP3, HELIOS and CD25 from 4 independent donors in 4 independent experiments. Paired t-test was used to determine statistical significance of the differences observed. p values are stated.

To directly examine the role of TNFR2 in human Treg proliferation, we deleted *TNFR2* gene in purified Tregs using the CRISPR/Cas9 technology. On day 4 after activation, only 27.3% of aCD3/CD28 bead stimulated cells electroporated with gRNA targeting the *TNFR2 gene* were TNFR2^hi^ when compared to 87.75% of cells that were electroporated with scrambled gRNA (Supplemental Figure 4A). Similarly, 14.5% of CD28SA stimulated cells were TNFR2^hi^ when compared to 52.6% of cells that were electroporated with scrambled gRNA (Supplemental Figure 4B), indicating successful TNFR2 deletion in the majority of the Tregs. We observed that TNFR2KO Tregs have severely and significantly reduced ex-vivo expansion with either aCD3/CD28 bead or CD28SA stimulation when compared with control Tregs (Figure 5B), demonstrating a positive role of TNFR2 signaling in ex-vivo proliferation of human Tregs. It is worth noting that, on day 8 post activation, 44.2% of aCD3/CD28 bead-stimulated and 44.6% of CD28SA-stimulated cells were TNFR2^hi^, increased from those detected on day 4 after stimulation. This suggested that the few TNFR2-sufficient Tregs after CRISPR editing had a proliferative advantage over their TNFR2KO counterparts (Supplemental Figure 4A and 4B). Consistent with our observation using etanercept, TNFR2KO Tregs showed decreased expression of CD25, FOXP3, and HELIOS when compared to TNFR2^hi^ Tregs (Supplemental Figure 4C and 4D). Together, these results demonstrated a role of TNFR2 in promoting human Treg ex-vivo proliferation.

Finally, we exposed TNFR2KO Tregs to IL-6 and TNFα to assess the requirement for TNFR2 in Treg proliferative boost by these cytokines shown in Figure 1. We observed that TNFR2KO Tregs had a diminished proliferative response to IL-6 and TNFα when compared to control Tregs (Figure 5C). Moreover, TNFR2KO Tregs had significantly decreased expression of key Treg lineage markers CD25, FOXP3, and HELIOS (Figure 5D). Together, these results suggest that TNFR2 signaling enhanced human Treg proliferation and contributed to preserving Treg identity during exposure to proinflammatory cytokines.

### Beadless protocol for ex-vivo expansion of human Tregs

Ex-vivo expanded human Tregs are currently being evaluated in many early phase clinical trials in transplantation and autoimmune diseases (66). Most of the Treg manufacturing processes rely on multiple rounds of stimulation with aCD3/28 beads (67). Our results of highly efficient expansion of lineage stable human Tregs using one cycle of CD28SA stimulation in the presence of IL-2, IL-6, and TNFα prompted us to consider this protocol as an alternative approach to expand Tregs for clinical use. We thus compared rates of Treg expansion induced with 1 or 2 rounds of aCD3/28 bead stimulation versus those achieved with single round of CD28SA stimulation with or without the addition of IL-6 and TNFα. Tregs stimulated with aCD3/28 beads entered cell expansion more rapidly than that induced by CD28SA, but the cells rested by day 9 and required restimulation to proliferate again (Figure 6A). In contrast, Tregs stimulated with CD28SA continue to proliferate and began to rest by 14 days after stimulation. Addition of IL-6, TNFα, or both did not lead more rapid entry into cell division, but more persistent proliferation resulting in more total cell yields at the end of the two-week expansion (Figure 6A). Given the concern of the reported negative impact of IL-6 on Tregs, we determined if the persistent Treg proliferation can be achieved using less IL-6. Side-by-side comparison of Treg expansion using 3 unrelated donors showed altering IL-6 concentrations from 15 ng/ml to 150 ng/ml in the context of CD28SA stimulation in the presence of 300 IU/ml IL-2 and 50ng/ml TNFα resulted in comparable Treg expansion over 14 days (Figure 6B).

**Figure 6.**
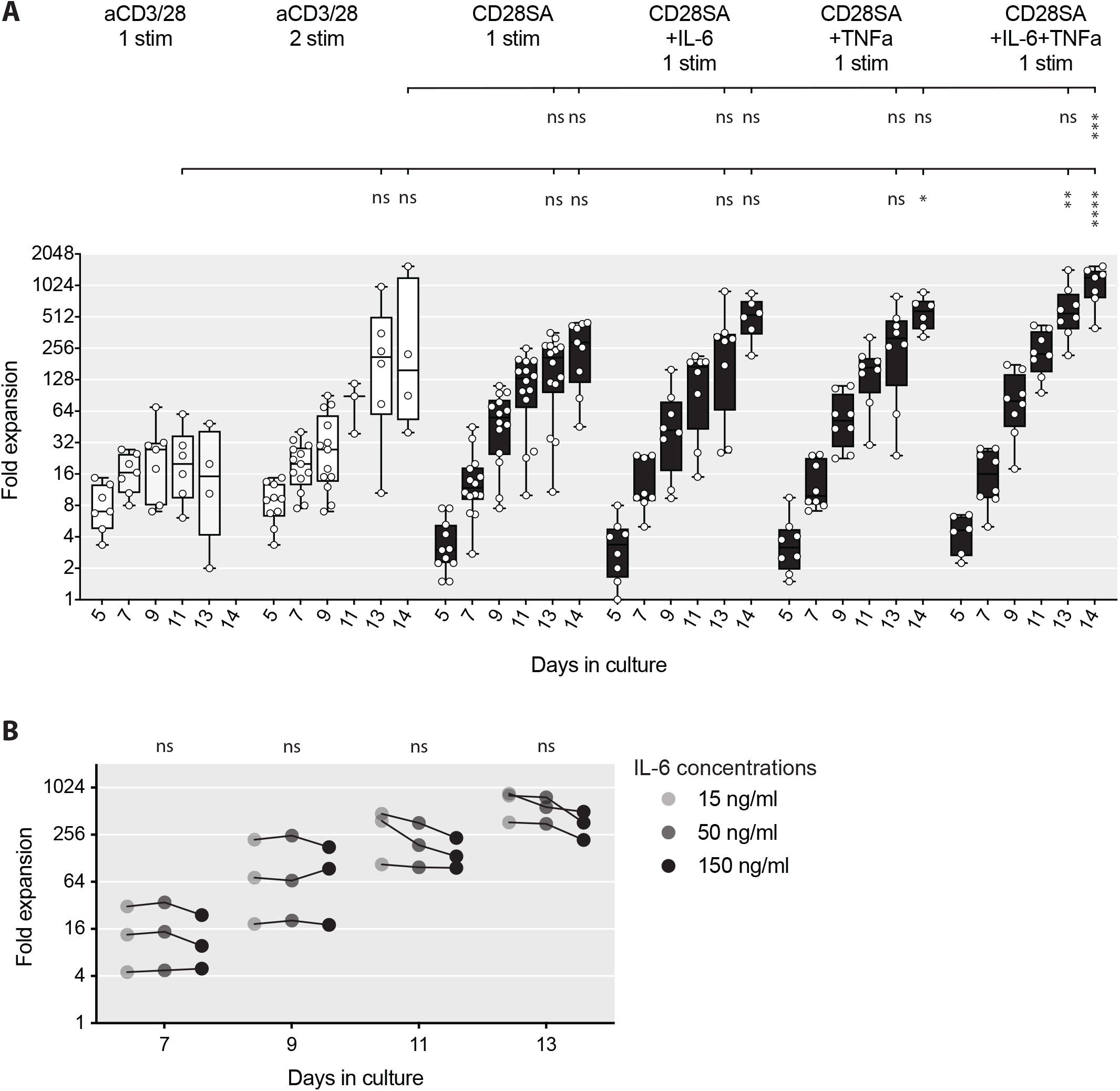
Beadless ex-vivo expansion of human Tregs. (**A**) FACS purified human Tregs were stimulated with either aCD3/28 beads or CD28SA and cultured in the presence of 300 IU/ml rhIL-2 with or without 150 ng/ml IL-6 and/or 50 ng/ml TNFa for up to 14 days. Expansion kinetics of Tregs from 7 to 14 unrelated donors in independent experiments are shown. Statistical significance of the differences was assessed using one-way ANOVA and Dunnett’s multiple comparisons posttest using aCD3/28 1 stim Day 13 and aCD3/28 2 stim Day 14 as baseline references. (**B**) FACS purified human Tregs were stimulated with CD28SA and cultured in the presence of 300 IU/mL rhIL-2, 50 ng/ml TNFa and varying concentrations of IL-6 (15 ng/ml, 50 ng/ml, and 150 ng/ml) as indicated. Expansion kinetics over 14 days of Tregs of from 3 unrelated donors in 3 independent experiments are shown. Data connected by the same line are from the same donor. Statistical significance of the differences was assessed using one-way ANOVA with Geisser-Greenhouse’s correction and Dunnett’s multiple comparisons posttest using 150 ng/ml IL-6 as baseline reference. p values are marked as *=p<0.05, **=p<0.01, ***=p<0.001, ****=p<0.0001.

The pattern of more persistent proliferation after CD28SA stimulation in the presence of TNFα and IL-6 when compared with aCD3/28 bead stimulation suggested that Tregs cultured in these two protocols were in distinct metabolic state. We thus performed metabolomic profiling of aCD3/28 bead-stimulated versus CD28SA+IL-6+TNFα beadless-stimulated Tregs. We selected day 7 for the comparison because the Tregs in both protocols were briskly proliferating and had comparable fold expansion at that time, but about to diverge in the rate of proliferation. We also included freshly purified Tregs as baseline. In total, 9 samples from 3 unrelated donors were profiled with targeted quantitative analysis using capillary electrophoresis mass spectrometry. Intracellular concentrations of 116 metabolites involved in glycolysis, pentose phosphate pathway, TCA cycle, lipid metabolism, urea cycle, and polyamine, creatine, purine, glutathione, nicotinamide, choline, and amino acid metabolisms were captured (Supplemental Table 1). The results for each of the 116 metabolites were graphed and superimposed on the pathway map of overall energy metabolism. Tregs on day 7 after activation with beads or the beadless protocol had significantly higher concentrations of all 20 amino acids than Tregs freshly purified from the peripheral blood (Figure 7A, Supplemental Table 1). All the amino acids, except aspartate, were present at equal or higher concentration in Tregs expanded with the beardless protocol when compared to those expanded with aCD3/28 beads. It was also notable that cysteine was only detected in Tregs stimulated with the beadless protocol and not detected in fresh and bead-stimulated Tregs. Both aspartate and cysteine are non-essential amino acids, and their altered levels may indicate altered metabolism of these amino acids in Tregs under different culture conditions.

**Figure 7.**
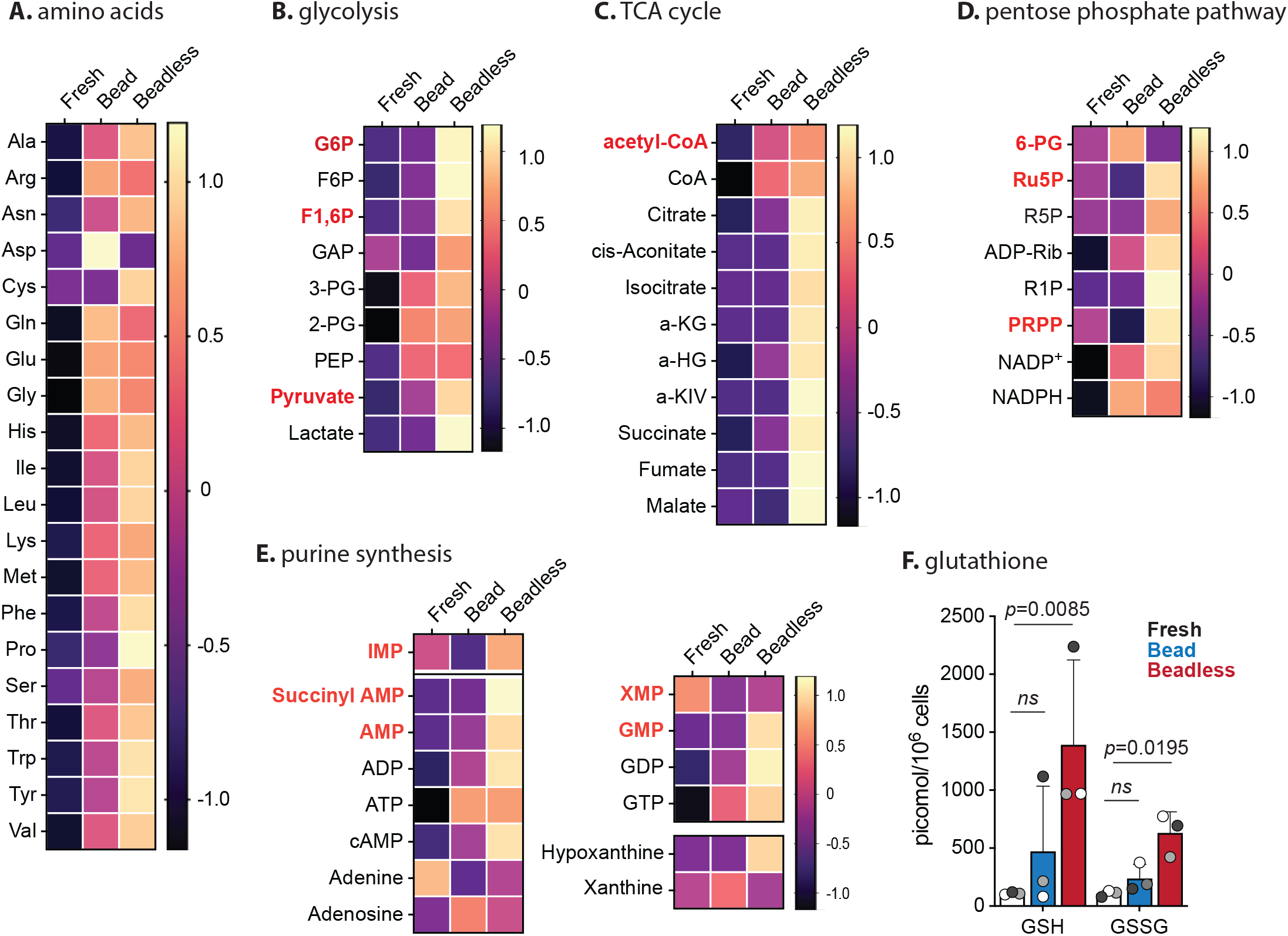
Metabolomic profile of Tregs before and after bead or beadless activation. Tregs were purified from 3 unrelated healthy donors. Tregs from each donor were divided into three parts: fresh Tregs without in vitro stimulation, Tregs expanded with aCD3/28 beads and IL-2 for 7 days (Bead), and Tregs expanded with CD28SA, IL-6, TNFa, and IL-2 for 7 days (Beadless). Intracellular metabolites were extracted and subjected to capillary electrophoresis mass spectrometry to profile 116 metabolites. The amount of each metabolite was normalized to the cell number and expressed as pmol/106 cells (Supplemental Table 1). The data for each metabolite were then normalized to the mean of all the samples and log2 transformed so that differences among the three experimental conditions can be compared across different metabolites. Non-detected metabolites were given a value of 2-52. The transformed data for amino acids (**A**), glycolysis (**B**), TCA cycle (**C**), pentose phosphate pathway (**D**), and purine synthesis (**E**) are summarized as heatmaps. Products of rate-limiting steps or key metabolites are highlighted with bold red text. (**F**) Intracellular concentration of reduced (GSH) and oxidized (GSSG) glutathione are shown. Circles represent individual data and data from the same Tregs donor are represented by the same fill color. Statistical significance of the differences was determined using ratio paired t test. p values are as marked. ns=not significant, p>0.05.

Glycolysis was activated by both protocols with a trend of higher concentrations of glycolytic intermediates in the Tregs stimulated with the beadless protocol (Figure 7B). Notably, the Tregs in the beadless protocol contained 10 times higher concentration of lactic acid when compared to the bead stimulated Tregs, indicating highly active glycolytic activities in these cells. Concurrently, the beadless protocol stimulated Tregs also had higher oxidative phosphorylation (OXPHOS) activities indicated by the higher concentrations of TCA intermediates when compared to bead stimulated Tregs, which were nearly completely depleted of TCA intermediates of cis-aconitate, isocitrate, alpha-ketoglutarate, fumarate and malate (Figure 7C, Supplemental Table 1). Thus, Tregs in the beadless protocol were in a high energy state with concurrent activation of both glycolysis and OXPHOS whereas bead stimulated Tregs mostly relied on glycolysis for energy production.

The pentose phosphate pathway also utilizes glycolysis intermediates to generate phosphoribosyl diphosphate, which serves as a substrate for purine and pyrimidine synthesis. Intermediates in the pentose phosphate pathway were elevated in the beadless protocol expanded Tregs when compared to fresh and bead stimulated Tregs (Figure 7D). This was correlated with higher concentrations of intermediates of purine biosynthesis (Figure 7E). The pentose phosphate pathway also converts NADP+ to NADPH, thus an important regulator of the redox state of the cell. Moreover, NADPH also supports de novo fatty acid synthesis. NADP+ and NADPH were present at comparable elevated concentrations in bead versus beadless stimulated Tregs (Figure 7D) whereas the NADPH to NADP+ ratios were both comparably reduced from the fresh Treg baseline (Supplemental Table 1), suggesting higher antioxidant demand and/or higher fatty acid synthesis in the activated Tregs. NADPH protects mitochondria against oxidative stress by transferring its reductive power to oxidized glutathione disulfide (GSSG) to generate reduced glutathione (GSH). GSH and GSSG concentrations were consistently low in the fresh Tregs from three donors, variably and moderately increased in bead stimulated Tregs, and significantly increased in Tregs activated with beadless stimulations (Figure 7F). Tregs activated with the beadless protocol had higher levels of total glutathione, suggestive of greater buffering ability for elevated intracellular reactive oxygen species (ROS). The increased energy production and de novo nucleotide synthesis combined with higher antioxidant potential may underscore the better metabolic fitness and more persistent proliferation of Tregs stimulated with the beadless protocol when compared to those stimulated with aCD3/28 beads.

### TNFα and IL-6-exposed Tregs maintained their function in vitro and in vivo

Lastly, we compared the function of Tregs expanded using aCD3/28 beads versus the beadless protocol. For the in vitro suppression assay, we used aCD3/28 beads expanded CD4^+^CD25^-^ CD127^hi^ Tconv as responders and mixed in titrated numbers of Tregs expanded with either aCD3/28 beads or CD28SA, with or without IL-6 and TNFα. Tregs expanded with various protocols all exhibited similar suppressive function, suggesting that ex-vivo exposure to IL-6 and TNFα did not negatively affect Treg suppressive function (Figure 8A).

**Figure 8.**
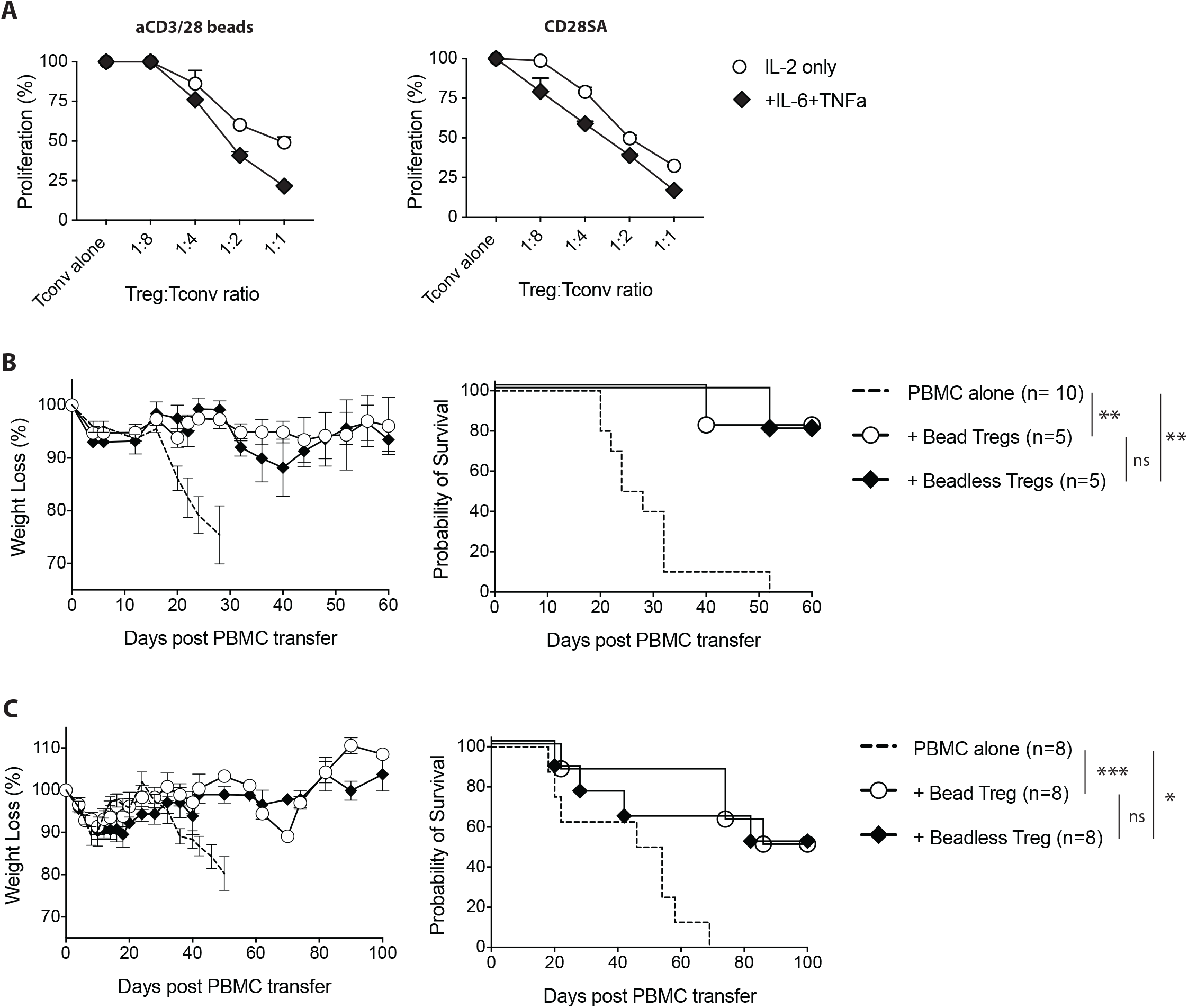
Tregs expanded with the beadless protocol maintained their in-vitro and in-vivo suppressive function. (**A**) FACS purified human Tregs were stimulated with either aCD3/28 beads or CD28SA and cultured in the presence of 300 IU/ml rhIL-2 with or without TNFa and/or IL-6 for 9 days, as indicated. In addition, FACS purified human Tconvs were stimulated with aCD3/28 beads and cultured in parallel to autologous Tregs. At day 7 of culture, expanded Tregs were harvested and then co-cultured with 5x 104 ex-vivo expanded Tconv at Treg/ Tconv ratios of 1:1, 1:2, 1:4 and 1:8 in the presence of aCD3/CD28 beads (1:10 bead to Tconv ratio). Proliferation was measured using 3[H]-thymidine incorporation in triplicate wells on day 4 of co-culture. Results shown are representative of 2 independent experiments using 2 unrelated donors. (**B**) At day 9 of culture, either standard protocol (aCD3/28 beads and IL-2) or beadless protocol (CD28SA, IL-2, IL-6 and TNFa) expanded Tregs were adoptively co-transferred into sub lethally irradiated NSG mice along with autologous PBMC. Results shown are from 1 experiment with a total of at least 5 mice per group. (**C**) Same as panel B, except the Tregs were injected 4 days after the PBMC infusion. Weight loss and survival were monitored every other day post-adoptive transfer. Results shown are sum-mary of 2 independent experiments using 2 unrelated donors with a total of 8 mice per group. Kaplan-Mei-er survival graphs are shown, and a log-rank comparison of the groups was used to calculate p values. p values are marked as ns=not significant, p>0.05, *=p<0.05, **=p<0.01, ***=p<0.001, ****=p<0.0001.

To measure the in-vivo suppressive function of Tregs, we used models of xenogeneic GVHD in the NSG mice. Intravenous injection of 5 × 10^6^ PBMC’s into sub lethally irradiated NSG mice resulted in the development GVHD that was lethal in 20 to 30 days (Figure 8B). Co-transfer of 5 × 10^6^ Tregs expanded with standard aCD3/28 bead protocol or the beadless protocol significantly attenuated weight loss and improved survival from GVHD in most of the mice (Figure 8B). To compare the function of the Tregs expanded with the standard versus beadless protocols in a more challenging condition, we infused Treg 4 days after PBMC injection. It has been previously reported that co-infusion of Treg with GVHD-inducing PBMCs effectively prevents Tconv proliferation and improves survival, while delayed Treg infusion requires higher doses of Tregs to reverse disease that has already been initiated (68). Indeed, an increased percentages of humanized NSG mice treated with late Treg infusion eventually succumbed to GVHD compared to the GVHD prevention model with PBMC and Treg co-infusions. In this suboptimal model, there were no differences identified between Tregs expanded with the standard protocol versus the beadless protocol in their ability to prevent weight loss and reduce mortality (Figure 8C). In summary, these results demonstrated that ex-vivo expanded Tregs exposed to IL-6 and TNFα maintained their function in vitro and in vivo in humanized mouse models of GVHD.

## Discussion

In this study, we challenged purified peripheral blood human Tregs ex-vivo with IL-6 and/or TNFα to determine their direct effect on Tregs. Our data demonstrated that Tregs responded to TNFα and IL-6 by increasing their proliferation while maintaining their lineage identity and functional capacity. In fact, persistent proliferation and high FOXP3 expression of activated Tregs cultured in the absence of exogenous TNFα or IL-6 depended on TNFR2, suggesting autocrine activation of this pathway during Treg activation. Together these data show that human Tregs positively respond to proinflammatory cytokines.

The ability of Tregs to respond and dampen inflammation is essential for the resolution of immune responses and restoring normal immune homeostasis. Results from this study are consistent with reports in mice on the important function of TNFα-TNFR2 axis in Treg function at site of inflammation in models of colitis, GVHD and EAE (10, 32-34, 36). We further show that human Tregs engage this pathway even in the absence of exogenously provided TNFα. In addition to promoting Treg proliferation to support the numerical dominance of Tregs at the site of inflammation, TNFα may also support the preservation of Treg lineage identity by sustaining high expression of FOXP3 and CD25. Moreover, it has been shown previously that mouse Tregs produce CCL3 and CCL4 to attract activated T cells to suppress them (69). Results from this study further show that human Treg production of CCL3, CCL4, and CCL5 are enhanced by TNFα, suggesting that this function may be more important for Treg function at sites of inflammation. Although activated Tregs engage TNFR2 pathway to promote persistent proliferation and sustain high expression of FOXP3, exogenous provision of TNFα further boosted this effect, suggesting the endogenously produced TNFα and LTα does not maximally activate this pathway. Consistent with this notion, we noted that CD28SA induced less TNFα (Figure 4A), which may partially explain the greater effect of exogenous TNFα on CD28SA stimulated Tregs.

At the start of our study, we noted a dominant theme of a negative role of IL-6 on Tregs in the literature. The finding of a lack of destabilizing effect of IL-6 on human Tregs despite the use of supraphysiological concentration of IL-6 and reduced IL-2 is thus unexpected. A further review of the literature shows a nearly unanimous conclusion that IL-6 antagonizes in vitro induction of Tregs from Tconvs by driving the cells to differentiation into Th17 effectors (22). Moreover, there is consensus that IL-6 enhances Teff activation and instill the ability to resist suppression by Tregs (20, 23). The body of literature on a direct role of IL-6 on lineage committed Tregs, especially on human Tregs, is comparatively more limited. An uncertainty when working with human Tregs is the identity of the cells at the start of the experiments depending on the markers used and gating strategy for Treg isolation. In this study, we isolated human Tregs from peripheral blood using FACS with markers of CD4^+^CD25^+^CD127^lo/-^. The addition of CD127 allows isolation of human Tregs with higher purity and higher yield (70, 71). Isolation of human Tregs based on the expression of CD4 and CD25 using magnetic activated cell sorting often results in less pure Tregs and outgrowth of Tconvs that necessitates the additions of TGFβ or rapamycin in Treg expansion cultures to preserve uniformly high FOXP3 expression (72-74). Even when the same markers are used, the placement of the CD4^+^CD25^+^CD127^lo/-^ gate requires operators’ judgement thus can introduce variations. Since neither CD25 expression nor CD127 downregulation is unique to Tregs and recently activated T cells can also express varying amounts of these markers, different gates can result in differences in the purity of the isolated Tregs. In this study, we used conservative gating to ensure the isolation of highly pure Tregs before proinflammatory cytokine challenge. In addition, Tregs are heterogenous thus different gate placements may lead to enrichment or depletion of certain subsets of Tregs that may respond differently to antigen and cytokine stimulations (75). Whether subsets of human Tregs respond differently to proinflammatory cytokines requires further investigation.

In this study, we noted significantly more robust proliferation of Tregs stimulated with CD28SA when compared to aCD3/28 bead stimulation. Furthermore, IL-6 and TNFα synergistically enhanced the proliferation stimulated by CD28SA. Notably, aCD3/28 bead stimulated faster entry into cell cycle but earlier rest akin to logarithmic growth; whereas CD28SA stimulated Tregs showed slower start but more robust and persistent cell cycling as seen in exponential growth, especially when IL-6 and TNFα are added. This suggests that Tregs are capable of exponential growth and the pattern of expansion stimulated by aCD3/28 beads may be a result of negative feedback that limits proliferation. The biochemical basis of the distinct proliferation patterns remains to be elucidated. In Tconvs, TCR stimulation in the absence of CD28 signaling activate nuclear factor of activated T-cells that is not balanced with concurrent activation of NF-κB and activator protein 1, which leads induction of E3 ubiquitin ligases Cbl-b, cell cycle arrest, and cellular anergy (76). CD28-mediated costimulation complement TCR signaling by activating NF-κB and AP1 while releasing the cells from the inhibitory effect of Cbl-b (76). We speculate that stimulation of Tregs with strong TCR agonist such as bead-bound aCD3/28 induces imbalanced activation of transcription factors that triggers negative feedback to limit Treg proliferation, resulting in short bursts of proliferation followed by stagnation (77).

On the other hand, CD28SA stimulated T cell activation depends on the expression of TCR but does not ostensibly induce activation of signaling intermediates immediately downstream of the TCR such as CD3 chains and ZAP70 (78, 79). Thus, CD28SA delivers a strong costimulatory signal in the context of a weak TCR signal (80-82), which may avoid the engagement of negative feedback loops that leads to earlier cell cycle arrest. The addition of IL-6 and TNFa may further engage complementary signaling pathways and transcription factors to boost Treg proliferation. Ongoing work in our laboratory is actively testing these predications.

Another non-mutually exclusive explanation for the distinct Treg proliferation patterns is the divergent metabolic programs induced by aCD3/28 bead versus CD28SA plus IL-6 and TNFα (beadless protocol). Tregs, like Tconv, undergo metabolic reprogramming after activation to accommodate the anabolic and energy demands for cell proliferation (83). Previous studies have reported either preferential fatty acid oxidation (FAO)-fueled oxidative phosphorylation over glycolysis by Tregs (84-86) or reliance on both glycolysis and FAO to support Treg proliferation and suppressive function (87, 88). In our metabolomic analysis, both bead and beadless stimulations led to increased intracellular amino acid concentrations, energy production, and nucleotide synthesis when compared to freshly isolated Tregs, consistent with their anabolic state at day 7 after stimulation. We noted three major distinctions in the metabolic state of Tregs stimulated with aCD3/28 beads versus the beadless protocol. First, while both protocols increased glycolytic flux, the beadless protocol Tregs had a slightly higher concentration of pyruvate, but 10 times more intracellular lactate. The conversion of pyruvate to lactate is coupled with oxidation of NADH to replenish the NAD+ pool in the cytosol, which is essential in preventing stagnation of glycolysis from NAD+ shortage (89). This suggest that Tregs in the beadless protocol had more active glycolysis. Second, the beadless protocol stimulated Tregs had higher concentrations of all TCA intermediates measured, whereas intermediates downstream of citrate were almost completely depleted in aCD3/28 bead stimulated Tregs. This suggests that the aCD3/28 bead stimulated Tregs have very low level of OXPHOS and relied mostly on glycolysis for energy production. Third, concentrations of total and reduced glutathione were consistently higher in the beadless protocol stimulated Tregs, suggestive of greater buffering capability for ROS. Intense OXPHOS can increase the generation of ROS leading to cell death. Therefore, reduced glutathione and other antioxidants are essential for maintaining cells in a high energetic state by scavenging ROS (90). GSH deficiency in Tregs results in an imbalanced intracellular redox state and impaired suppressive function (91). Overall, our analysis of Treg intracellular metabolites indicates that the beadless protocol stimulated Tregs have higher glycolytic flux, more activate oxidative phosphorylation, and higher antioxidation potential that may contribute to their persistent proliferation.

It is unclear at this point of our investigation which component of the beadless protocol contributed to the more favorable metabolic program in Tregs. CD28 costimulation is known to be required for inducing glycolysis in activated T cells during proliferation (92, 93), and can also enhance the spare respiratory capacity in T cells in a dose dependent manner, priming mitochondria for increased OXPHOS demand (94). CD28SA has been shown to drive T effector memory cells into an adaptable metabolic state that can flexibly maximize glycolysis and OXPHOS potentials depending on glucose and oxygen availability (95), but the impact of CD28SA on metabolic programming of Tregs remains uncertain. Another component of the beadless protocol, TNFα may act through TNFR2, to induces a glycolytic switch in Tregs coupled with shunting of intermediates into the TCA cycle, thereby promoting anabolic biosynthetic processes (96). Lastly, it has been reported that some STAT3 activated by IL-6 can localize in the inner membrane of mitochondria and enhance the efficiency of electron transport chain and reduce the generation of ROS (97). Further research is needed to dissect the role of the individual components of the beadless protocol on Treg metabolism.

A practical implication of the findings of this study is a new protocol for in vitro expansion of human Tregs. Given the pivotal role of Tregs in immune system homeostasis, Treg therapy has emerged as a potential immunoregulatory therapy in transplantation and autoimmune diseases. One of the challenges currently facing Treg therapy is the ability to reliably manufacture enough Tregs in a short period of time without the need for repeated stimulations that negatively affect Treg stability (98). The beadless protocol described in this study can promote highly efficient and consistent expansion of human Tregs with only one cycle of CD28SA stimulation. Another advantage of the CD28SA-based protocol is the use of all soluble reagents, thus harvesting the Treg products at the end of the expansion is simplified without the need to remove the stimulating beads. Importantly, Tregs manufactured with the beadless protocol have demethylated FOXP3 enhancer and are suppressive in vitro and in vivo in humanized mouse GVHD models. Thus, the beadless protocol offers several improvements in the Treg manufacturing process compared to the currently used standard protocol.

Taken together, our findings show that human Tregs positively respond to TNFα and IL-6 by increased proliferation without compromising their lineage stability. With proper stimulation and the right cytokine milieu, human Tregs can grow exponentially, which may be a result of balanced transcription and metabolic programing. We speculate that increased proliferation in response to inflammatory cytokines allows Tregs to scale to inflammation to restore immune homeostasis. These properties of Tregs may be harnessed to improve the manufacture of therapeutic Tregs for autoimmune diseases and transplantation.

## Materials and methods

### Human peripheral blood samples

De-identified peripheral blood units were purchased from StemCell Technologies (Vancouver, Canada). Peripheral blood samples from 25 healthy donors (11 male and 14 female) between the age of 21 and 72 were used.

### Ex-vivo Treg culture

Human blood samples were processed the same day after collection using ficoll (Cytiva, Marlborough MA) density gradient to isolate PBMC. PBMC were stained with anti-CD4 FITC (clone OKT4), anti-CD25 APC (clone 4E3) and anti-CD127 PE (clone HIL-7R-M21, all from BD Biosciences, San Jose, CA). CD4^+^CD25^+^CD127^lo/-^ Tregs and CD4^+^CD25^-^CD127^+^ Tconvs were purified using FACS. Post sort analyses confirmed >99% purity. Purified Tregs were plated in 48-well plate at a density of 1×10^5^ cells/well in medium consisted of 10% heat-inactivated fetal bovine serum (Biosource International), nonessential amino acids, 0.5 mM sodium pyruvate, 5 mM Hepes and 1 mM glutaMax I (all from Invitrogen) in an RPMI-1640 base. Human anti-CD3 and anti-CD28-conjugated dynabeads (aCD3/CD28 beads; Fischer Scientific, Hampton, NH) were added to the T cells at 1:1 bead to cell ratio. Alternatively, CD28 superagonist (CD28SA) (5μg/mL; clone anti-CD28.1; Ancell, Stillwater, MN) were used to stimulate the T cells.

Recombinant human IL-2 (Proleukin, Prometheus laboratories, Switzerland) were added at concentration indicated. Recombinant human IL-6 (50ng/ml; Peprotech, Rocky Hill, NJ) and recombinant human TNFα (50ng/ml; Peprotech, Rocky Hill, NJ) were also added in selected wells. Etanercept (5μg/mL; ETN-Enbrel, Pfizer) was added in selected experiments. Ex-vivo expansion cultures were supplemented with fresh media containing IL-2 on day 2, 5, and every other day thereafter, until the cells returned quiescent state (typically, day 9 for aCD3/CD28 bead stimulated Tregs and day 14 for CD28SA stimulated Tregs). CD4^+^ Tconvs were similarly expanded prior to use for subsequent experiments.

### Flow cytometry

Single-cell suspensions were recovered from ex-vivo expanded Treg cultures. The cellular suspensions were first stained with surface antibodies to CD4 (Per-CP; clone SK3), CD25 (PE-Cy7; clone 4E3), TNFR2 (APC; clone hTNFR-M1), and fixable viability dye (APC-e780, all from purchased from BD Biosciences) prior to fixation and permeabilization using kit according to manufacturer protocol (catalog number 88-8824-00; Thermofisher) and intracellular staining for FOXP3 (e450; clone PCH101) and HELIOS (FITC; clone 22F6, both from BD Biosciences). The samples were analyzed on a FACS LSR-II flow cytometer (BD Biosciences) and analyzed using FACSDiva (BD Biosciences) or Flow Jo software (Tree Star, Ashland, OR).

### In-vitro Treg suppression assay

Ex-vivo expanded human Tregs were harvested on day 7 of culture and varying numbers of Tregs were then co-cultured with 1× 10^5^ ex-vivo expanded Tconv at Treg/ Tconv ratios of 1:1, 1:2, 1:4 and 1:8 in the presence of aCD3/CD28 beads (1:10 bead to Tconv ratio). 0.5μCi [^3^H]-Thymidine was added 16hrs before the cells were harvested for analysis on day 4 of co-culture. Triplicate wells were established for each condition.

### TSDR analysis

Ex-vivo expanded Tregs (2×10^6^ from each condition) were harvested on day 7 of culture and then cryopreserved in FBS containing 10% DMSO at −80°C. The samples were submitted to Epigendx (Boston, MA) for analysis of DNA methylation of the TSDR in FOXP3 intron 1 using the ADS783FS2 assay.

### Cytokine assay

Supernatants of Treg and Tconv cultures were aspirated on day 7 and sent for analysis using a 42-plexed Luminex human cytokine detection kit (EveTechnologies, Vancouver, Canada).

### Humanized NSG mouse graft-versus-host disease (GVHD) models

Eight to twelve-week-old NOD.Cg-Prkdc^scid^Il2rg^tm1Wjl^/SzJ (NSG, Jackson Laboratory, Stock No 005557) male mice were bred in our animal facilities in specific pathogen-free conditions. To induce GVHD, NSG mice were irradiated (2.5Gy) one day prior to retroorbital vein infusion of either 4×10^6^ fresh PBMCs or ex-vivo expanded Tconvs. In GVHD prevention experiments, some mice receive co-infusion of ex-vivo expanded Tregs. In GVHD treatment experiments, Tregs were infused 4 days after PBMC infusion. Xenogeneic GVHD development was evaluated by clinical examination and body weight measurements for either 60 days (prevention experiments) or 100 days (treatment experiments).

### *TNFR2* gene deletion using CRISPR

Ribonucleoprotein complexes were made by complexing crRNAs and tracrRNAs chemically synthesized (Integrated DNA Technologies (IDT), Coralville, IA) with recombinant Cas9NLS (QB3 Macrolab, UC Berkley, CA) as previously described(99). Guide RNA sequences used for gene editing were (1) GGTTCTTGACTACCGTAATT (scrambled gRNA) and (2) GGCAUUUACACCCUACGCCC (*TNFR2* gene). Briefly, Lyophilized RNAs were resuspended at 160μM in 10mM Tris-HCL with 150mM KCL and stored in aliquots at minus 80. The day of electroporation, CrRNA and tracrRNA aliquots were thawed and mixed at a 1:1 volume and incubated 30 minutes at 37°C for annealing. The 80µM guide RNA complex was mixed at 37°C with Cas9 NLs at a 2:1 gRNA to Cas9 molar ratio for another 15 minutes and the mixture is defined as RNP at 20µM. RNP were electroporated using a Lonza 4D 96-well electroporation system (code EH115). Genome editing efficiency was measured by assessing cell surface TNFR2 expression using flow cytometry on days 4 and 8 of culture.

### Metabolomics

At least 2× 10^6^ FACS sorted fresh and ex-vivo expanded Tregs were collected on days 0 and 7 of culture, respectively. The cells were stored according to manufacturer protocol prior to being sent for analysis using capillary electrophoresis- and liquid chromatography-mass spectrometry platforms to generate their global metabolic profile (Human Metabolome Technologies, Boston, MA).

### Statistics

Kaplan-Meier survival graphs were constructed, and a log-rank comparison of the groups was used to calculate p-values. The paired t test was used for comparison of experimental groups. Differences were considered significant for p less than 0.05. Prism software (GraphPad Software, San Diego, CA) version 9 was used for data analysis and graphing data.

### Study approval

All experiments were approved (IACUC protocol No. AN183959-02) and conducted in accordance with UCSF IACUC regulations.

## Abbreviations

aCD3/28: anti-CD3 and anti-CD28 dynabeads
CD28SA: CD28 super agonist
CNS: Central Nervous System
FAO: Fatty acid oxidation
FOXP3: Forkhead box protein
P3 GVHD: Graft-versus-host-disease
GSH: glutathione
GSSG: glutathione disulfide
LTα: lymphotoxin α
NSG: NOD.Cg-Prkdc^scid^Il2rg^tm1Wjl^/SzJ
OXPHOS: oxidative phosphorylation
RORγT: Retinoic acid-related orphan receptor γT
Tconv: conventional CD4^+^ T cell
Teff: effector T cell
TNFR1: Tumor necrosis factor receptor 1
TNFR2: Tumor necrosis factor receptor 2
TNFSF: Tumor necrosis factor superfamily
TCR: T cell receptor
TSDR: Treg-specific demethylated region

## Author contributions

Designed the project: QT, NS. Supervised the project: QT, FV. Designed experiments: NS, QT. Performed experiments: NS, YP, LMRF, VN. Analyzed data: NS, YP, QT. Provided reagents and advice: YDM, FV. Wrote the manuscript: NS, QT.

## Acknowledgements

This work was supported by grants from NIAID/CTOT (Clinical Trials in Organ Transplant, 0255-B011-4609 and 0255-B012-4609, which were ancillary to a NIAID grant U01 AI113362), The Leona M. and Harry B. Helmsley Charitable Trust (2018PG-T1D042), NIDDK (UC4 DK116264). This works used the UCSF Flow Cytometry Core, which is supported in part by the Diabetes Research Center Grant from NIDDK (P30 DK063720 and P30 DK063720). NS was supported by a fellowship grant from the American Society of Nephrology. YDM was supported by the Swiss National Science Foundation (Advanced Postdoctoral Mobility Grant no.

P300PB_174500) and a fellowship grant from the University Hospital of Geneva. LMRF was the Jeffrey G. Klein Family Diabetes Fellow. We would like to thank Juan Du for the technical support and lab management. We would also like to thank the Marson lab for technical support. Finally, we would like to thank Jeffrey Bluestone for critical review of this manuscript.

## References

1. Sakaguchi, S., et al., Regulatory T cells and immune tolerance. Cell, 2008. 133(5): p. 775–87.

2. Rudensky, A.Y., Regulatory T cells and Foxp3. Immunol Rev, 2011. 241(1): p. 260–8.

3. Tang, Q. and J.A. Bluestone, The Foxp3+ regulatory T cell: a jack of all trades, master of regulation. Nat Immunol, 2008. 9(3): p. 239–44.

4. Vignali, D.A., Mechanisms of T(reg) Suppression: Still a Long Way to Go. Front Immunol, 2012. 3: p. 191.

5. Smigiel, K.S., et al., Regulatory T-cell homeostasis: steady-state maintenance and modulation during inflammation. Immunol Rev, 2014. 259(1): p. 40–59.

6. D’Alessio, F.R., et al., CD4+CD25+Foxp3+ Tregs resolve experimental lung injury in mice and are present in humans with acute lung injury. J Clin Invest, 2009. 119(10): p. 2898–913.

7. D’Alessio, F.R., et al., Lung Angiogenesis Requires CD4(+) Forkhead Homeobox Protein-3(+) Regulatory T Cells. Am J Respir Cell Mol Biol, 2015. 52(5): p. 603–10.

8. D’Alessio, F.R., J.T. Kurzhagen, and H. Rabb, Reparative T lymphocytes in organ injury. J Clin Invest, 2019. 129(7): p. 2608–2618.

9. Spence, A., et al., Revealing the specificity of regulatory T cells in murine autoimmune diabetes. Proc Natl Acad Sci U S A, 2018. 115(20): p. 5265–5270.

10. Ronin, E., et al., Tissue-restricted control of established central nervous system autoimmunity by TNF receptor 2-expressing Treg cells. Proc Natl Acad Sci U S A, 2021. 118(13).

11. Zhang, N., et al., Regulatory T cells sequentially migrate from inflamed tissues to draining lymph nodes to suppress the alloimmune response. Immunity, 2009. 30(3): p. 458–69.

12. Luan, D., et al., FOXP3 mRNA Profile Prognostic of Acute T Cell-mediated Rejection and Human Kidney Allograft Survival. Transplantation, 2021. 105(8): p. 1825–1839.

13. Shang, B., et al., Prognostic value of tumor-infiltrating FoxP3+ regulatory T cells in cancers: a systematic review and meta-analysis. Sci Rep, 2015. 5: p. 15179.

14. Sakaguchi, S., et al., The plasticity and stability of regulatory T cells. Nat Rev Immunol, 2013. 13(6): p. 461–7.

15. Papadakis, K.A. and S.R. Targan, Tumor necrosis factor: biology and therapeutic inhibitors. Gastroenterology, 2000. 119(4): p. 1148–57.

16. Grivennikov, S.I., et al., Distinct and nonredundant in vivo functions of TNF produced by t cells and macrophages/neutrophils: protective and deleterious effects. Immunity, 2005. 22(1): p. 93–104.

17. Apostolaki, M., et al., Cellular mechanisms of TNF function in models of inflammation and autoimmunity. Curr Dir Autoimmun, 2010. 11: p. 1–26.

18. Tanaka, T., M. Narazaki, and T. Kishimoto, IL-6 in inflammation, immunity, and disease. Cold Spring Harb Perspect Biol, 2014. 6(10): p. a016295.

19. Yokota, S., T.D. Geppert, and P.E. Lipsky, Enhancement of antigen- and mitogen-induced human T lymphocyte proliferation by tumor necrosis factor-alpha. J Immunol, 1988. 140(2): p. 531–6.

20. Li, B., L.L. Jones, and T.L. Geiger, IL-6 Promotes T Cell Proliferation and Expansion under Inflammatory Conditions in Association with Low-Level RORgammat Expression. J Immunol, 2018. 201(10): p. 2934–2946.

21. Chen, X. and J.J. Oppenheim, Contrasting effects of TNF and anti-TNF on the activation of effector T cells and regulatory T cells in autoimmunity. FEBS Lett, 2011. 585(23): p. 3611–8.

22. Chen, X., O.M. Howard, and J.J. Oppenheim, Pertussis toxin by inducing IL-6 promotes the generation of IL-17-producing CD4 cells. J Immunol, 2007. 178(10): p. 6123–9.

23. Pasare, C. and R. Medzhitov, Toll pathway-dependent blockade of CD4+CD25+ T cell-mediated suppression by dendritic cells. Science, 2003. 299(5609): p. 1033–6.

24. Mercadante, E.R. and U.M. Lorenz, Breaking Free of Control: How Conventional T Cells Overcome Regulatory T Cell Suppression. Front Immunol, 2016. 7: p. 193.

25. Wohlfert, E.A. and R.B. Clark, ‘Vive la Resistance!’--the PI3K-Akt pathway can determine target sensitivity to regulatory T cell suppression. Trends Immunol, 2007. 28(4): p. 154–60.

26. Zhang, Q., et al., TNF-alpha impairs differentiation and function of TGF-beta-induced Treg cells in autoimmune diseases through Akt and Smad3 signaling pathway. J Mol Cell Biol, 2013. 5(2): p. 85–98.

27. Housley, W.J., et al., Natural but not inducible regulatory T cells require TNF-alpha signaling for in vivo function. J Immunol, 2011. 186(12): p. 6779–87.

28. Ni, X., et al., TRAF6 directs FOXP3 localization and facilitates regulatory T-cell function through K63-linked ubiquitination. EMBO J, 2019. 38(9).

29. Grinberg-Bleyer, Y., et al., Pathogenic T cells have a paradoxical protective effect in murine autoimmune diabetes by boosting Tregs. J Clin Invest, 2010. 120(12): p. 4558–68.

30. Hamano, R., et al., TNF optimally activatives regulatory T cells by inducing TNF receptor superfamily members TNFR2, 4-1BB and OX40. Eur J Immunol, 2011. 41(7): p. 2010–20.

31. Chen, X., et al., Interaction of TNF with TNF receptor type 2 promotes expansion and function of mouse CD4+CD25+ T regulatory cells. J Immunol, 2007. 179(1): p. 154–61.

32. Chen, X., et al., TNFR2 is critical for the stabilization of the CD4+Foxp3+ regulatory T. cell phenotype in the inflammatory environment. J Immunol, 2013. 190(3): p. 1076–84.

33. Atretkhany, K.N., et al., Intrinsic TNFR2 signaling in T regulatory cells provides protection in CNS autoimmunity. Proc Natl Acad Sci U S A, 2018. 115(51): p. 13051–13056.

34. Chen, X., et al., Cutting edge: expression of TNFR2 defines a maximally suppressive subset of mouse CD4+CD25+FoxP3+ T regulatory cells: applicability to tumor-infiltrating T regulatory cells. J Immunol, 2008. 180(10): p. 6467–71.

35. Pierini, A., et al., TNF-alpha priming enhances CD4+FoxP3+ regulatory T-cell suppressive function in murine GVHD prevention and treatment. Blood, 2016. 128(6): p. 866–71.

36. Leclerc, M., et al., Control of GVHD by regulatory T cells depends on TNF produced by T cells and TNFR2 expressed by regulatory T cells. Blood, 2016. 128(12): p. 1651–9.

37. Nie, H., et al., Phosphorylation of FOXP3 controls regulatory T cell function and is inhibited by TNF-alpha in rheumatoid arthritis. Nat Med, 2013. 19(3): p. 322–8.

38. Valencia, X., et al., TNF downmodulates the function of human CD4+CD25hi T-regulatory cells. Blood, 2006. 108(1): p. 253–61.

39. Stoop, J.N., et al., Tumor necrosis factor alpha inhibits the suppressive effect of regulatory T cells on the hepatitis B virus-specific immune response. Hepatology, 2007. 46(3): p. 699–705.

40. Nagar, M., et al., TNF activates a NF-kappaB-regulated cellular program in human CD45RA-regulatory T cells that modulates their suppressive function. J Immunol, 2010. 184(7): p. 3570–81.

41. Zaragoza, B., et al., Suppressive activity of human regulatory T cells is maintained in the presence of TNF. Nat Med, 2016. 22(1): p. 16–7.

42. Chen, X., et al., Co-expression of TNFR2 and CD25 identifies more of the functional CD4+FOXP3+ regulatory T cells in human peripheral blood. Eur J Immunol, 2010. 40(4): p. 1099–106.

43. Wang, J., et al., TNFR2 ligation in human T regulatory cells enhances IL2-induced cell proliferation through the non-canonical NF-kappaB pathway. Sci Rep, 2018. 8(1): p. 12079.

44. Okubo, Y., et al., Homogeneous expansion of human T-regulatory cells via tumor necrosis factor receptor 2. Sci Rep, 2013. 3: p. 3153.

45. He, X., et al., A TNFR2-Agonist Facilitates High Purity Expansion of Human Low Purity Treg Cells. PLoS One, 2016. 11(5): p. e0156311.

46. Brown, G., et al., Tumor necrosis factor-alpha inhibitor-induced psoriasis: Systematic review of clinical features, histopathological findings, and management experience. J Am Acad Dermatol, 2017. 76(2): p. 334–341.

47. Kemanetzoglou, E. and E. Andreadou, CNS Demyelination with TNF-alpha Blockers. Curr Neurol Neurosci Rep, 2017. 17(4): p. 36.

48. Korn, T., et al., IL-6 controls Th17 immunity in vivo by inhibiting the conversion of conventional T cells into Foxp3+ regulatory T cells. Proc Natl Acad Sci U S A, 2008. 105(47): p. 18460–5.

49. Xu, L., et al., Cutting edge: regulatory T cells induce CD4+CD25-Foxp3-T cells or are self-induced to become Th17 cells in the absence of exogenous TGF-beta. J Immunol, 2007. 178(11): p. 6725–9.

50. O’Connor, R.A., et al., Foxp3(+) Treg cells in the inflamed CNS are insensitive to IL-6-driven IL-17 production. Eur J Immunol, 2012. 42(5): p. 1174–9.

51. van Loosdregt, J., et al., Stabilization of the transcription factor Foxp3 by the deubiquitinase USP7 increases Treg-cell-suppressive capacity. Immunity, 2013. 39(2): p. 259–71.

52. Gao, Y., et al., Inflammation negatively regulates FOXP3 and regulatory T-cell function via DBC1. Proc Natl Acad Sci U S A, 2015. 112(25): p. E3246–54.

53. Fujimoto, M., et al., The influence of excessive IL-6 production in vivo on the development and function of Foxp3+ regulatory T cells. J Immunol, 2011. 186(1): p. 32–40.

54. Hagenstein, J., et al., A Novel Role for IL-6 Receptor Classic Signaling: Induction of RORgammat(+)Foxp3(+) Tregs with Enhanced Suppressive Capacity. J Am Soc Nephrol, 2019. 30(8): p. 1439–1453.

55. Samanta, A., et al., TGF-beta and IL-6 signals modulate chromatin binding and promoter occupancy by acetylated FOXP3. Proc Natl Acad Sci U S A, 2008. 105(37): p. 14023–7.

56. Lu, L., et al., Critical role of all-trans retinoic acid in stabilizing human natural regulatory T cells under inflammatory conditions. Proc Natl Acad Sci U S A, 2014. 111(33): p. E3432–40.

57. Bin Dhuban, K., et al., Signaling Through gp130 Compromises Suppressive Function in Human FOXP3(+) Regulatory T Cells. Front Immunol, 2019. 10: p. 1532.

58. Ferreira, R.C., et al., Human IL-6R(hi)TIGIT(-) CD4(+)CD127(low)CD25(+) T cells display potent in vitro suppressive capacity and a distinct Th17 profile. Clin Immunol, 2017. 179: p. 25–39.

59. Putnam, A.L., et al., Expansion of human regulatory T-cells from patients with type 1 diabetes. Diabetes, 2009. 58(3): p. 652–62.

60. He, X., et al., Single CD28 stimulation induces stable and polyclonal expansion of human regulatory T cells. Sci Rep, 2017. 7: p. 43003.

61. Gerriets, V.A., et al., Foxp3 and Toll-like receptor signaling balance Treg cell anabolic metabolism for suppression. Nat Immunol, 2016. 17(12): p. 1459–1466.

62. Kim, H.J., et al., Stable inhibitory activity of regulatory T cells requires the transcription factor Helios. Science, 2015. 350(6258): p. 334–9.

63. Yang, X.O., et al., Molecular antagonism and plasticity of regulatory and inflammatory T cell programs. Immunity, 2008. 29(1): p. 44–56.

64. Li, L., J. Kim, and V.A. Boussiotis, IL-1beta-mediated signals preferentially drive conversion of regulatory T cells but not conventional T cells into IL-17-producing cells. J Immunol, 2010. 185(7): p. 4148–53.

65. Locksley, R.M., N. Killeen, and M.J. Lenardo, The TNF and TNF receptor superfamilies: integrating mammalian biology. Cell, 2001. 104(4): p. 487–501.

66. Esensten, J.H., et al., Regulatory T-cell therapy for autoimmune and autoinflammatory diseases: The next frontier. J Allergy Clin Immunol, 2018. 142(6): p. 1710–1718.

67. Tang, Q. and J.A. Bluestone, Regulatory T-cell therapy in transplantation: moving to the clinic. Cold Spring Harb Perspect Med, 2013. 3(11).

68. Nguyen, V.H., et al., In vivo dynamics of regulatory T-cell trafficking and survival predict effective strategies to control graft-versus-host disease following allogeneic transplantation. Blood, 2007. 109(6): p. 2649–56.

69. Patterson, S.J., et al., T regulatory cell chemokine production mediates pathogenic T cell attraction and suppression. J Clin Invest, 2016. 126(3): p. 1039–51.

70. Liu, W., et al., CD127 expression inversely correlates with FoxP3 and suppressive function of human CD4+ T reg cells. J Exp Med, 2006. 203(7): p. 1701–11.

71. Seddiki, N., et al., Expression of interleukin (IL)-2 and IL-7 receptors discriminates between human regulatory and activated T cells. J Exp Med, 2006. 203(7): p. 1693–700.

72. Xia, G., J. He, and J.R. Leventhal, Ex vivo-expanded natural CD4+CD25+ regulatory T cells synergize with host T-cell depletion to promote long-term survival of allografts. Am J Transplant, 2008. 8(2): p. 298–306.

73. Fraser, H., et al., A Rapamycin-Based GMP-Compatible Process for the Isolation and Expansion of Regulatory T Cells for Clinical Trials. Mol Ther Methods Clin Dev, 2018. 8: p. 198–209.

74. Mathew, J.M., et al., A Phase I Clinical Trial with Ex Vivo Expanded Recipient Regulatory T cells in Living Donor Kidney Transplants. Sci Rep, 2018. 8(1): p. 7428.

75. Miyara, M., et al., Functional delineation and differentiation dynamics of human CD4+ T cells expressing the FoxP3 transcription factor. Immunity, 2009. 30(6): p. 899–911.

76. Paolino, M. and J.M. Penninger, Cbl-b in T-cell activation. Semin Immunopathol, 2010. 32(2): p. 137–48.

77. Hippen, K.L., et al., Massive ex vivo expansion of human natural regulatory T cells (T(regs)) with minimal loss of in vivo functional activity. Sci Transl Med, 2011. 3(83): p. 83ra41.

78. Tacke, M., et al., CD28-mediated induction of proliferation in resting T cells in vitro and in vivo without engagement of the T cell receptor: evidence for functionally distinct forms of CD28. Eur J Immunol, 1997. 27(1): p. 239–47.

79. Hunig, T. and K. Dennehy, CD28 superagonists: mode of action and therapeutic potential. Immunol Lett, 2005. 100(1): p. 21–8.

80. Bischof, A., et al., Autonomous induction of proliferation, JNK and NF-alphaB activation in primary resting T cells by mobilized CD28. Eur J Immunol, 2000. 30(3): p. 876–82.

81. Waibler, Z., et al., Signaling signatures and functional properties of anti-human CD28 superagonistic antibodies. PLoS One, 2008. 3(3): p. e1708.

82. Dennehy, K.M., et al., Mitogenic signals through CD28 activate the protein kinase Ctheta-NF-kappaB pathway in primary peripheral T cells. Int Immunol, 2003. 15(5): p. 655–63.

83. Pearce, E.L., et al., Fueling immunity: insights into metabolism and lymphocyte function. Science, 2013. 342(6155): p. 1242454.

84. Michalek, R.D., et al., Cutting edge: distinct glycolytic and lipid oxidative metabolic programs are essential for effector and regulatory CD4+ T cell subsets. J Immunol, 2011. 186(6): p. 3299–303.

85. Angelin, A., et al., Foxp3 Reprograms T Cell Metabolism to Function in Low-Glucose, High-Lactate Environments. Cell Metab, 2017. 25(6): p. 1282–1293 e7.

86. Howie, D., et al., Foxp3 drives oxidative phosphorylation and protection from lipotoxicity. JCI Insight, 2017. 2(3): p. e89160.

87. Procaccini, C., et al., The Proteomic Landscape of Human Ex Vivo Regulatory and Conventional T Cells Reveals Specific Metabolic Requirements. Immunity, 2016. 44(2): p. 406–21.

88. Cluxton, D., et al., Differential Regulation of Human Treg and Th17 Cells by Fatty Acid Synthesis and Glycolysis. Front Immunol, 2019. 10: p. 115.

89. Liberti, M.V. and J.W. Locasale, The Warburg Effect: How Does it Benefit Cancer Cells? Trends Biochem Sci, 2016. 41(3): p. 211–218.

90. Sena, L.A. and N.S. Chandel, Physiological roles of mitochondrial reactive oxygen species. Mol Cell, 2012. 48(2): p. 158–67.

91. Kurniawan, H., et al., Glutathione Restricts Serine Metabolism to Preserve Regulatory T Cell Function. Cell Metab, 2020. 31(5): p. 920–936 e7.

92. Frauwirth, K.A., et al., The CD28 signaling pathway regulates glucose metabolism. Immunity, 2002. 16(6): p. 769–77.

93. Jacobs, S.R., et al., Glucose uptake is limiting in T cell activation and requires CD28-mediated Akt-dependent and independent pathways. J Immunol, 2008. 180(7): p. 4476–86.

94. Klein Geltink, R.I., et al., Mitochondrial Priming by CD28. Cell, 2017. 171(2): p. 385–397 e11.

95. Thaventhiran, T., et al., CD28 Superagonistic Activation of T Cells Induces a Tumor Cell-Like Metabolic Program. Monoclon Antib Immunodiagn Immunother, 2019. 38(2): p. 60–69.

96. de Kivit, S., et al., Stable human regulatory T cells switch to glycolysis following TNF receptor 2 costimulation. Nat Metab, 2020. 2(10): p. 1046–1061.

97. Rincon, M. and F.V. Pereira, A New Perspective: Mitochondrial Stat3 as a Regulator for Lymphocyte Function. Int J Mol Sci, 2018. 19(6).

98. Hoffmann, P., et al., Loss of FOXP3 expression in natural human CD4+CD25+ regulatory T cells upon repetitive in vitro stimulation. Eur J Immunol, 2009. 39(4): p. 1088–97.

99. Roth, T.L., et al., Reprogramming human T cell function and specificity with non-viral genome targeting. Nature, 2018. 559(7714): p. 405–409.

