## Supplemental Figure for "TNFa and IL-6 promote ex-vivo proliferation of lineage-committed human regulatory T cells"

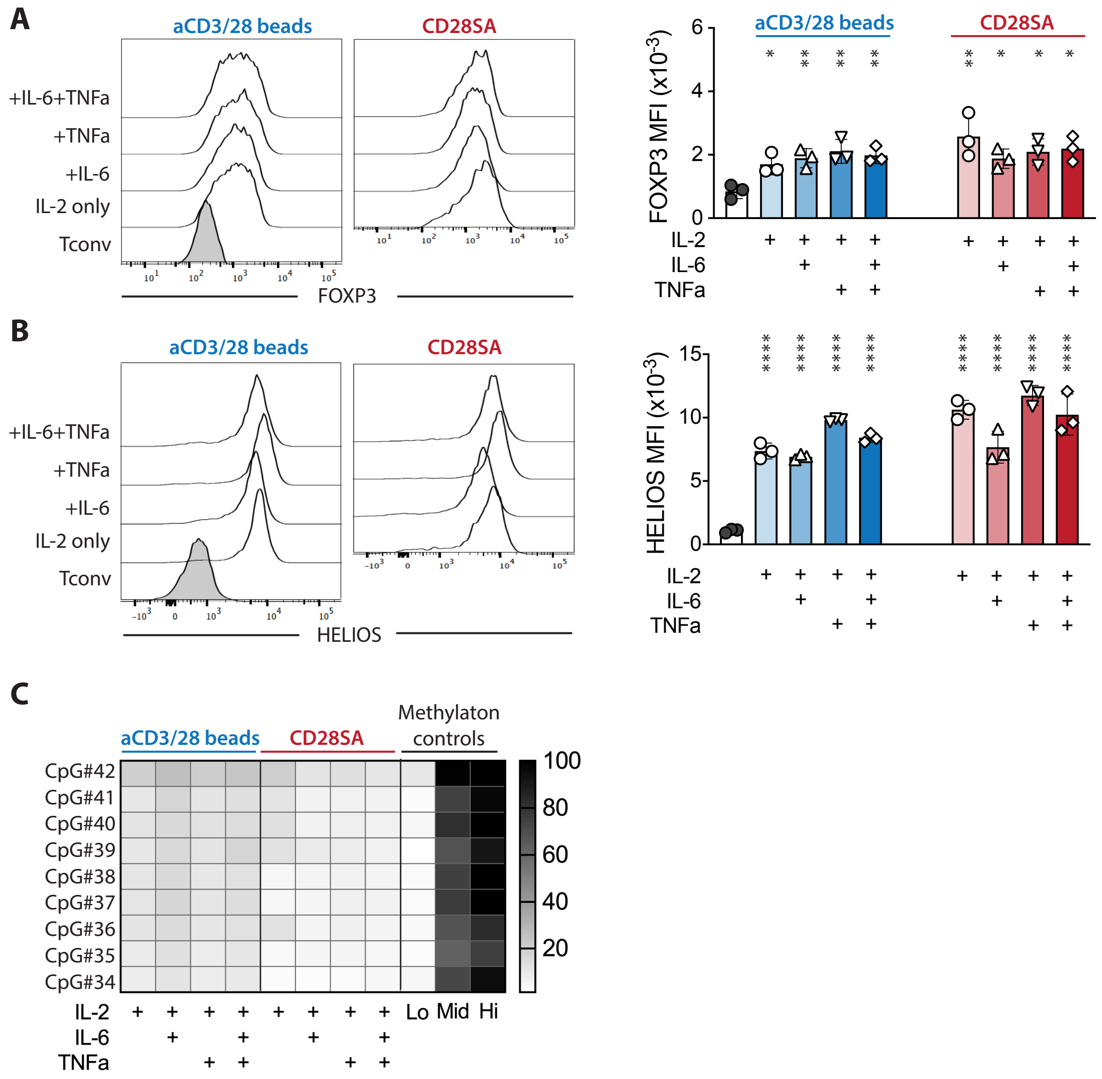


**Figure 1. Tregs exposed to IL-6 and/or TNFa maintained their lineage identity despite decreased IL-2.** FACS purified human Tregs were stimulated with either aCD3/28 beads or CD28SA and cultured in the presence of 15 IU/ml rhIL-2 with or without TNFa and IL-6 as indicated. (**A** and **B**) Flow cytometric analysis of FOXP3 (A) and HELIOS (B) expression in Tregs on day 9 after stimulation. Representative histograms (left) and summaries of MFI's from 3 independent experiments (right) are shown. Statistical significance of differences was assessed using one-way ANOVA and Dunnett's multiple comparisons test using Tconv (Panels A and B) as a baseline reference. p values are marked as *=p<0.05, **=p<0.01, ***=p<0.001, ****=p<0.0001. (**C**) Heatmap summary of TSDR demethylation of Tregs expanded in various conditions. Results shown are averages of Treg cultures using 2 unrelated male donors in 2 independent experiments.

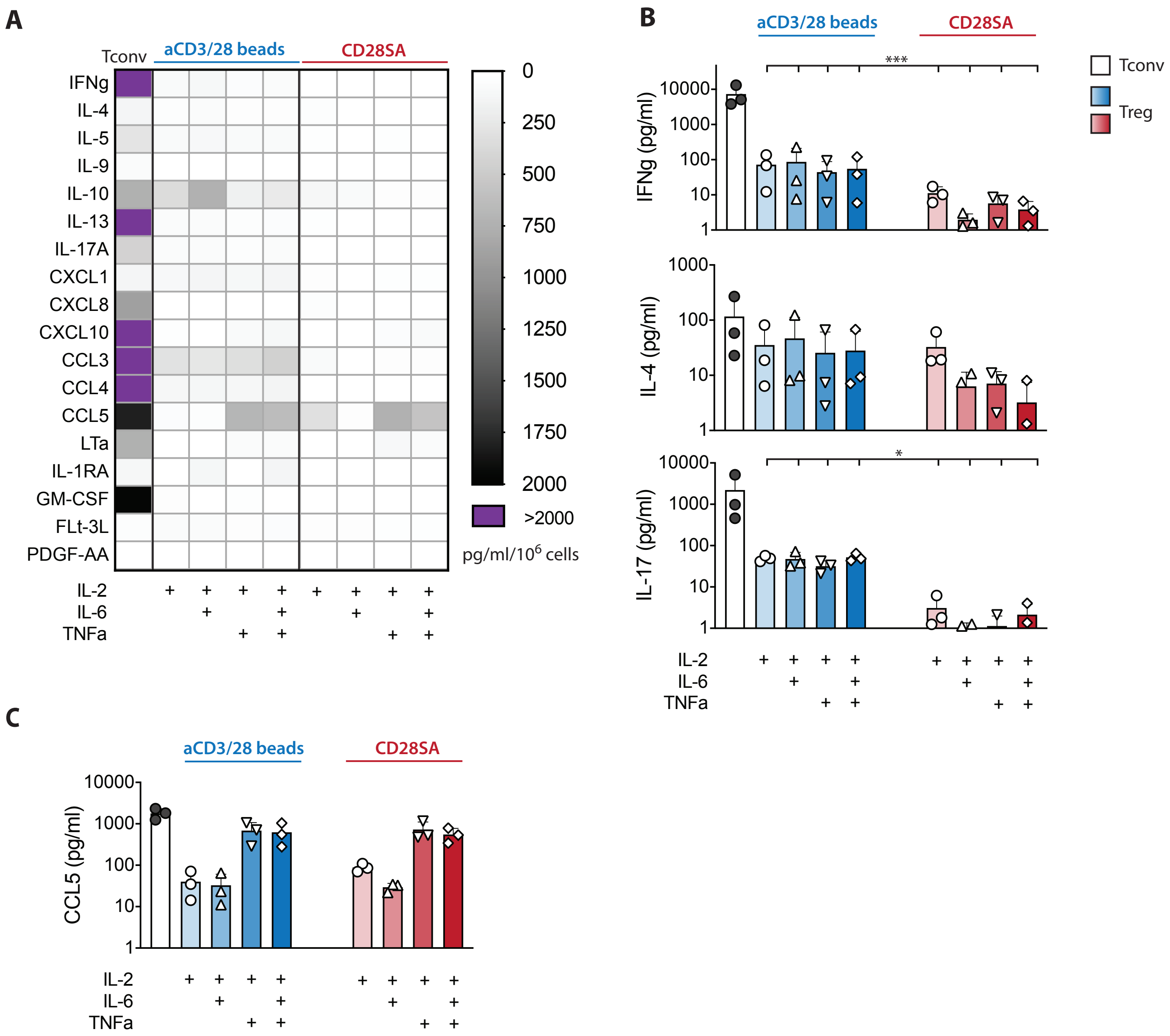


**Figure 2. TNFa and IL-6 expanded Tregs did not secrete proinflammatory cytokines despite decreased IL-2 challenge.** FACS purified human Tregs were stimulated with either aCD3/28 beads or CD28SA and cultured in the presence of 15 IU/ml rhIL-2 with or without TNFa and IL-6 as indicated. Cytokine and chemokine secretion in the culture supernatant of various Treg cultures was assessed using a multiplex Luminex panel. Supernatant in aCD3/28 bead stimulated Tconv cultures are included as a reference. (**A**) Heatmap summary of cytokines and chemokines that were present in any of the culture condition is shown. (**B**) IFN-g, IL-4, and IL-17 concentrations in the Day 7 culture supernatants are shown. (**C**) CCL5 and LTa concentrations in the Day 7 culture supernatant are shown. Results shown are summaries of 3 independent experiments using cells from 3 unrelated donors. Statistical significance of differences was assessed using one-way ANOVA and Dunnett's multiple comparisons posttest using Tconv (Panels B and C) as a baseline reference. p values are marked as *=p<0.05, **=p<0.01, ***=p<0.001, ****=p<0.0001.

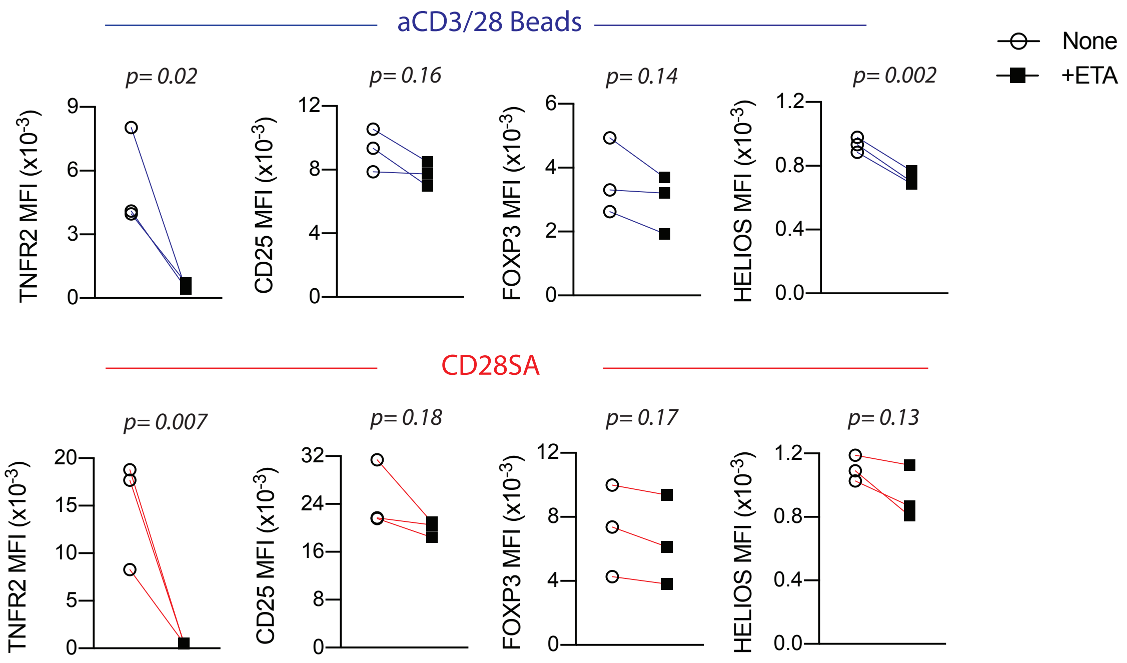


**Figure 3. Etanercept-treated Tregs have decreased TNFR2, CD25, FOXP3 and HELIOS expression.** Summary of MFI of TNFR2, CD25, FOXP3 and HELIOS expres-sion of Tregs cultured in the presence or absence of etanercept, assessed by flow cytometry at day 8 of culture. Paired t-test was used for statistical analysis. Results shown are summary of 3 independent donors from 3 independent experiments. p values are stated.

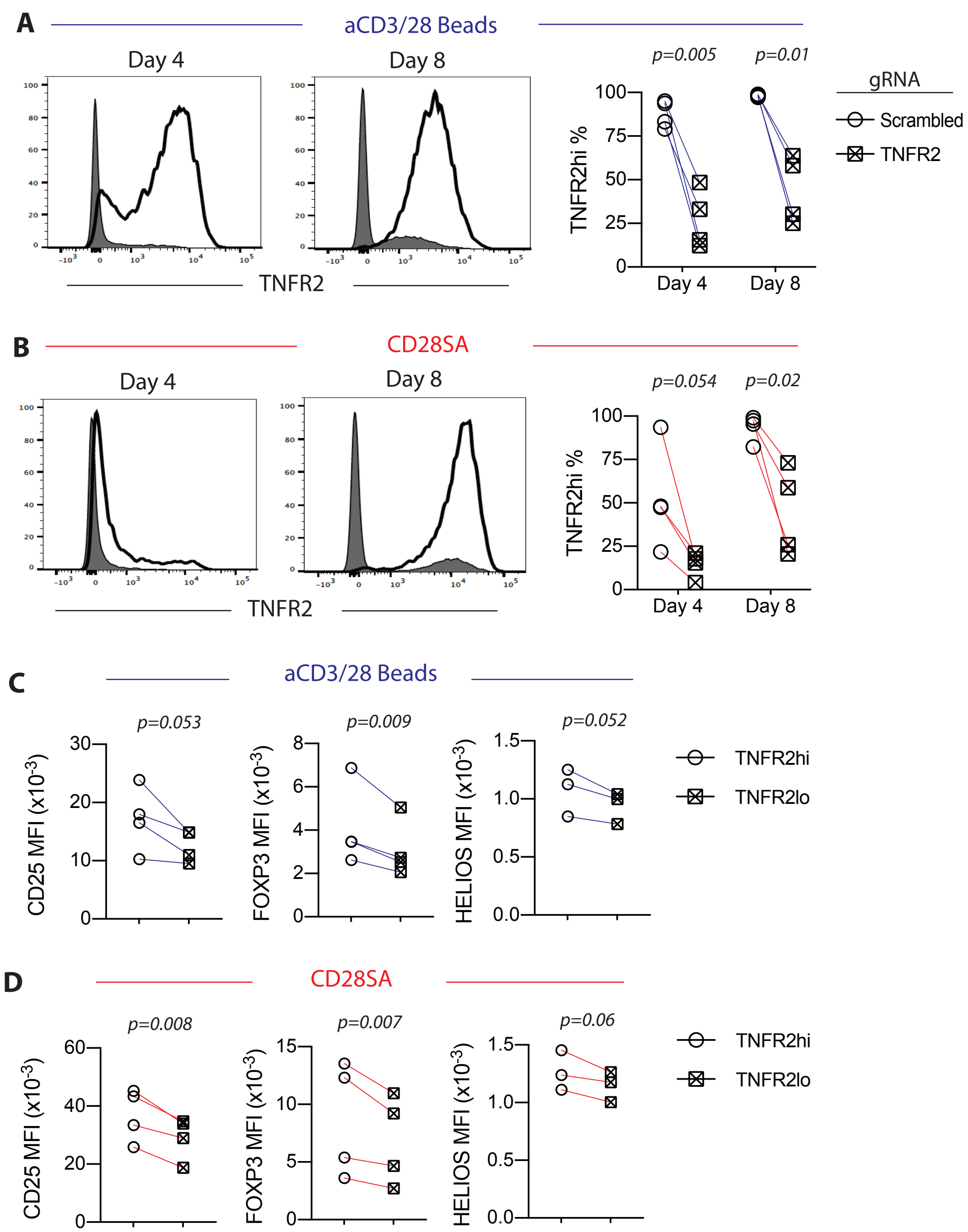


**Figure 4. TNFR2KO Tregs generated with CRISPR-Cas9 technology have decreased CD25, FOXP3 and HELIOS expression.** Human Tregs were gene edited using the CRISPR-Cas9 technology to delete the TNFR2 gene prior to stimulation with either anti-CD3/CD28 beads or anti-CD28SA. TNFR2 expression of either (**A**) aCD3/28 bead or (**B**) CD28SA stimulated Tregs that were electroporated with scrambled gRNA (open histograms) or gRNA against the TNFR2 gene (shaded histograms) was assessed by flow cytometry on day 4 and 8 of ex-vivo Treg culture. Paired t-test was used for statisti-cal analysis. MFI's of CD25, FOXP3 and HELIOS of either (**C**) aCD3/28 bead or (**D**) CD28SA stimulated TNFR2hi versus TNFR2KO Tregs were assessed by flow cytometry on day 8 of culture. Paired t-test was used for statistical analysis. Results shown are summary of 4 independent donors from 4 independent experiments. p values are stated.

### Absolute Concentration of 116 Compounds (pmol/$10^6$ cells)

| | Compound name | Pathway Label | PubChem CID | Fresh | | | Bead | | | Beadless | | |
|---|---|---|---|---|---|---|---|---|---|---|---|---|
| | | | | Donor 1 | Donor 2 | Donor 3 | Donor 1 | Donor 2 | Donor 3 | Donor 1 | Donor 2 | Donor 3 |
| A_0001 | NAD$^+$ | NAD+ | 5893 | 7.1 | 6.7 | 6.4 | 160 | 141 | 215 | 421 | 227 | 327 |
| A_0002 | cAMP | cAMP | 6076 | 0.2 | 0.3 | 0.3 | 0.5 | 0.5 | 0.4 | 1.4 | 0.7 | 0.7 |
| A_0003 | cGMP | cGMP | 24316 | N.D. | N.D. | N.D. | N.D. | N.D. | N.D. | N.D. | N.D. | N.D. |
| A_0004 | NADH | NADH | 439153 | 8.3 | 5.0 | 3.9 | 14 | 20 | 14 | 38 | 20 | 32 |
| A_0005 | Xanthine | Xanthine | 1188 | 1.1 | N.D. | 0.4 | 1.6 | N.D. | 0.6 | 0.5 | N.D. | 0.8 |
| A_0006 | ADP-ribose | ADP-Rib | 445794 | 0.05 | 0.04 | 0.03 | 0.6 | 0.4 | 1.0 | 1.5 | 0.6 | 1.5 |
| A_0007 | Mevalonic acid | Mevalonic acid | 134965 | N.D. | N.D. | N.D. | N.D. | N.D. | N.D. | N.D. | N.D. | N.D. |
| A_0008 | UDP-glucose | UDP-Glc | 8629 | 4.7 | 7.7 | 8.1 | 71 | 52 | 98 | 273 | 152 | 255 |
| A_0009 | Uric acid | Uric acid | 1175 | 3.3 | 6.2 | 1.4 | 1.6 | 1.1 | N.D. | 2.1 | 0.3 | N.D. |
| A_0010 | NADP$^+$ | NADP+ | 5886 | 2.0 | 0.9 | 0.8 | 3.1 | 4.1 | 4.4 | 5.9 | 4.3 | 5.3 |
| A_0011 | IMP | IMP | 8582 | 77 | 10 | 7.4 | 2.1 | 2.8 | 16 | 83 | 23 | 51 |
| A_0012 | Sedoheptulose 7-phosphate | S7P | 165007 | N.D. | N.D. | N.D. | N.D. | N.D. | N.D. | N.D. | N.D. | N.D. |
| A_0013 | Glucose 6-phosphate | G6P | 5958 | 3.8 | 8.1 | 4.3 | 8.6 | 7.9 | 2.9 | 13 | 14 | 17 |
| A_0014 | Fructose 6-phosphate | F6P | 603 | 1.4 | 2.8 | 1.5 | 3.5 | 3.0 | 1.8 | 5.3 | 6.3 | 7.6 |
| A_0015 | Fructose 1-phosphate | D-F1P | 439394 | N.D. | N.D. | N.D. | N.D. | N.D. | N.D. | 14 | 7.9 | N.D. |
| A_0016 | Galactose 1-phosphate | Gal1P | 123912 | N.D. | N.D. | N.D. | N.D. | N.D. | N.D. | 1.4 | 1.0 | 3.1 |
| A_0017 | Glucose 1-phosphate | G1P | 65533 | 1.7 | 1.2 | 1.0 | 3.2 | 3.7 | 1.8 | 3.2 | 2.4 | 4.9 |
| A_0018 | Acetoacetyl CoA | AAcCoA | 92153 | N.D. | N.D. | N.D. | N.D. | N.D. | N.D. | N.D. | N.D. | N.D. |
| A_0019 | Acetyl CoA | AcCoA | 444493 | 0.2 | 0.2 | 0.13 | 1.0 | 1.3 | 1.6 | 2.0 | 0.09 | 3.6 |
| A_0020 | Folic acid | Folic acid | 6037 | 0.3 | 0.3 | 0.3 | 1.2 | 1.5 | 0.7 | 0.4 | 0.2 | 0.6 |
| A_0021 | Ribose 5-phosphate | R5P | 439167 | 3.3 | 3.8 | 2.1 | 1.9 | 2.7 | 4.2 | 5.6 | 1.8 | 6.7 |
| A_0022 | CoA | CoA | 87642 | 0.8 | 0.6 | 0.4 | 1.7 | 1.8 | 3.2 | 2.6 | 2.4 | 2.9 |
| A_0023 | Ribose 1-phosphate | R1P | 439236 | 1.4 | 0.9 | 0.9 | 2.1 | N.D. | 1.6 | 7.0 | 3.4 | 4.8 |
| A_0024 | Ribulose 5-phosphate | Ru5P | 439184 | 0.4 | 0.8 | N.D. | 0.09 | N.D. | 0.3 | 1.0 | 0.5 | 1.4 |
| A_0025 | Xylulose 5-phosphate | X5P | 439190 | N.D. | N.D. | N.D. | N.D. | N.D. | N.D. | N.D. | N.D. | N.D. |
| A_0026 | Erythrose 4-phosphate | E4P | 122357 | N.D. | N.D. | N.D. | N.D. | N.D. | N.D. | N.D. | N.D. | N.D. |
| A_0027 | HMG CoA | HMG-CoA | 445127 | 0.3 | 0.2 | 0.2 | 0.5 | 0.8 | 0.4 | 1.5 | 0.7 | 1.3 |
| A_0028 | Glyceraldehyde 3-phosphate | Glyceraldehyde 3-phosphate | 729 | 0.8 | 1.2 | 0.9 | N.D. | N.D. | 1.6 | 1.6 | 0.2 | 4.2 |
| A_0029 | NADPH | NADPH | 5884 | 6.5 | 4.0 | 3.3 | 10 | 14 | 8.2 | 12 | 7.8 | 11 |
| A_0030 | Malonyl CoA | Malonyl-CoA | 644066 | N.D. | N.D. | N.D. | 0.4 | 0.7 | 0.2 | 0.3 | 0.12 | 0.4 |
| A_0031 | Phosphocreatine | Phosphocreatine | 9548602 | 0.011 | 0.3 | 0.3 | 3.0 | N.D. | 0.05 | 2.8 | 0.12 | N.D. |
| A_0032 | XMP | XMP | 73323 | 4.2 | 0.2 | 0.14 | 0.3 | 0.3 | 0.2 | 0.9 | 0.3 | 0.6 |
| A_0033 | Dihydroxyacetone phosphate | DHAP | 668 | 3.9 | 3.1 | 0.7 | N.D. | N.D. | 1.8 | 1.4 | 2.0 | 4.6 |
| A_0034 | Adenylosuccinic acid | Succinyl AMP | 447145 | 1.7 | 0.7 | 0.4 | 1.0 | 1.5 | 1.2 | 6.5 | 3.3 | 5.7 |
| A_0035 | Fructose 1,6-diphosphate | F1,6P | 172313 | 7.3 | 30 | 18 | 27 | 26 | 18 | 43 | 59 | 34 |
| A_0036 | 6-Phosphogluconic acid | 6-PG | 91493 | 1.5 | 11 | 7.9 | 21 | 9.7 | 4.8 | 6.7 | 3.6 | 5.3 |
| A_0037 | *N*-Carbamoylaspartic acid | Carbamoyl-Asp | 93072 | N.D. | 0.2 | 2.2 | 1.4 | 0.6 | 2.0 | 21 | 26 | 24 |
| A_0038 | PRPP | PRPP | 7339 | 8.8 | 34 | 48 | 4.7 | 6.3 | 21 | 72 | 53 | 52 |
| A_0039 | 2-Phosphoglyceric acid | 2-PG | 439278 | 0.4 | 0.5 | 0.5 | 1.2 | 0.7 | 1.0 | 1.0 | 1.0 | 1.1 |
| A_0040 | 2,3-Diphosphoglyceric acid | Diphosphoglycerate | 186004 | 1.3 | 5.9 | 6.8 | 9.6 | 6.3 | 17 | 16 | 26 | 16 |
| A_0041 | 3-Phosphoglyceric acid | 3-PG | 439183 | 3.5 | 3.7 | 4.1 | 8.5 | 4.3 | 8.4 | 8.5 | 8.2 | 8.3 |
| A_0042 | Phosphoenolpyruvic acid | PEP | 1005 | N.D. | 0.6 | 0.9 | 1.6 | N.D. | 2.3 | 1.9 | 0.9 | 1.3 |
| A_0043 | GMP | GMP | 6804 | 15 | 4.7 | 2.3 | 3.8 | 4.9 | 18 | 37 | 14 | 29 |
| A_0044 | AMP | AMP | 6083 | 38 | 8.6 | 5.1 | 7.8 | 7.5 | 105 | 177 | 58 | 145 |
| A_0045 | 2-Oxoisovaleric acid | 2-KIV | 49 | N.D. | N.D. | N.D. | N.D. | N.D. | N.D. | 5.5 | 4.0 | 6.6 |
| A_0046 | GDP | GDP | 8977 | 13 | 8.4 | 6.3 | 22 | 19 | 53 | 103 | 50 | 88 |
| A_0047 | Lactic acid | Lactic acid | 612 | 112 | 114 | 218 | 1,959 | 1,618 | 1,307 | 15,714 | 7,418 | 14,009 |
| A_0048 | ADP | ADP | 6022 | 56 | 30 | 27 | 100 | 88 | 335 | 489 | 254 | 449 |
| A_0049 | GTP | GTP | 6830 | 34 | 61 | 54 | 273 | 177 | 348 | 440 | 287 | 387 |
| A_0050 | Glyoxylic acid | Glyoxylic acid | 760 | N.D. | N.D. | N.D. | N.D. | N.D. | N.D. | N.D. | N.D. | N.D. |
| A_0051 | ATP | ATP | 5957 | 157 | 290 | 274 | 1,586 | 1,078 | 1,838 | 1,619 | 1,198 | 1,677 |
| A_0052 | Glycerol 3-phosphate | Glycerol 3-phosphate | 439162 | 4.1 | 4.3 | 8.9 | 126 | 103 | 219 | 310 | 134 | 178 |
| A_0053 | Glycolic acid | Glycolic acid | 757 | 50 | 23 | 19 | 51 | 115 | 24 | 23 | 14 | N.D. |
| A_0054 | Pyruvic acid | Pyruvic acid | 1060 | 36 | 22 | 32 | 83 | 75 | 55 | 140 | 76 | 230 |
| A_0055 | *N*-Acetylglutamic acid | N-AcGlu | 70914 | N.D. | 1.0 | 0.9 | 2.1 | N.D. | 2.9 | 1.8 | 2.0 | 3.1 |
| A_0056 | 2-Hydroxyglutaric acid | 2-Hydroxyglutaric acid | 43 | 1.3 | N.D. | 0.4 | 2.2 | 2.7 | 1.2 | 5.2 | 3.6 | 7.1 |
| A_0057 | Carbamoylphosphate | Carbamoyl-P | 278 | N.D. | N.D. | N.D. | N.D. | N.D. | 0.13 | N.D. | N.D. | N.D. |
| A_0058 | Succinic acid | Succinic acid | 1110 | 15 | 7.3 | 8.6 | 22 | 33 | 26 | 94 | 52 | 93 |
| A_0059 | Malic acid | Malic acid | 525 | 56 | 26 | 20 | N.D. | N.D. | 35 | 325 | 241 | 292 |
| A_0060 | 2-Oxoglutaric acid | 2-OG | 51 | N.D. | N.D. | N.D. | N.D. | N.D. | N.D. | 63 | 22 | 30 |
| A_0061 | Fumaric acid | Fumaric acid | 444972 | 1.6 | N.D. | 0.4 | N.D. | N.D. | 4.9 | 65 | 41 | 56 |
| A_0062 | Citric acid | Citric acid | 311 | 8.4 | 50 | 63 | 116 | 124 | 82 | 425 | 286 | 243 |
| A_0063 | *cis*-Aconitic acid | cis-Aconitic acid | 643757 | N.D. | 1.0 | 1.5 | 1.3 | 0.4 | 1.1 | 11 | 6.3 | 4.9 |
| A_0064 | Isocitric acid | Isocitric acid | 1198 | N.D. | N.D. | 0.06 | N.D. | N.D. | N.D. | 16 | 8.8 | 3.3 |
| C_0001 | Urea | Urea | 1176 | N.D. | N.D. | N.D. | N.D. | N.D. | N.D. | N.D. | N.D. | N.D. |
| C_0002 | Gly | Gly | 750 | 145 | 222 | 212 | 1,697 | 1,410 | 2,199 | 1,507 | 1,041 | 2,141 |
| C_0003 | Putrescine | Putrescine | 1045 | N.D. | N.D. | N.D. | N.D. | N.D. | N.D. | N.D. | N.D. | N.D. |
| C_0004 | Ala | Ala | 602 | 25 | 116 | 136 | 766 | 315 | 430 | 800 | 330 | 1,098 |
| C_0005 | β-Ala | b-Ala | 239 | 5.6 | 7.9 | 3.0 | 142 | 33 | 54 | 427 | 65 | 86 |
| C_0006 | Sarcosine | Sarcosine | 1088 | N.D. | N.D. | N.D. | N.D. | N.D. | N.D. | N.D. | N.D. | N.D. |
| C_0007 | γ-Aminobutyric acid | g-Aminobutyric acid | 119 | 15 | 8.4 | 3.1 | 12 | N.D. | 3.9 | 21 | 4.7 | 9.3 |
| C_0008 | *N*,*N*-Dimethylglycine | DMG | 673 | N.D. | N.D. | N.D. | N.D. | N.D. | N.D. | N.D. | N.D. | N.D. |
| C_0009 | Choline | Choline | 305 | 50 | 34 | 67 | 343 | 183 | 445 | 275 | 251 | 383 |
| C_0010 | Ser | Ser | 617 | 30 | 60 | 82 | 138 | 138 | 159 | 108 | 134 | 607 |
| C_0011 | Carnosine | Carnosine | 439224 | 1.1 | 2.7 | 1.4 | 0.3 | N.D. | 1.7 | 0.4 | N.D. | N.D. |
| C_0012 | Creatinine | Creatinine | 588 | 1.3 | 2.6 | 1.5 | 3.2 | N.D. | N.D. | 4.2 | N.D. | 0.7 |
| C_0013 | Pro | Pro | 614 | 42 | 129 | 138 | 358 | 144 | 842 | 1,783 | 956 | 1,904 |

| | Compound name | Pathway Label | PubChem CID | Fresh | | | Bead | | | Beadless | | |
|---|---|---|---|---|---|---|---|---|---|---|---|---|
| | | | | Donor 1 | Donor 2 | Donor 3 | Donor 1 | Donor 2 | Donor 3 | Donor 1 | Donor 2 | Donor 3 |
| C_0014 | Betaine | Betaine | 247 | N.D. | N.D. | N.D. | N.D. | N.D. | N.D. | N.D. | N.D. | N.D. |
| C_0015 | Val | Val | 1182 | N.D. | 37 | 52 | 219 | 229 | 351 | 314 | 286 | 637 |
| C_0016 | Thr | Thr | 6288 | 56 | 82 | 83 | 369 | 391 | 632 | 466 | 512 | 1,111 |
| C_0017 | Homoserine | Homoserine | 12647 | N.D. | N.D. | N.D. | N.D. | N.D. | N.D. | N.D. | N.D. | N.D. |
| C_0018 | Betaine aldehyde | BTL | 249 | N.D. | N.D. | N.D. | N.D. | N.D. | N.D. | N.D. | N.D. | N.D. |
| C_0019 | Cys | Cys | 594 | N.D. | N.D. | N.D. | N.D. | N.D. | N.D. | N.D. | 2.3 | 5.1 |
| C_0020 | Hydroxyproline | Hydroxyproline | 5810 | N.D. | 7.4 | 6.2 | 46 | 21 | N.D. | 35 | N.D. | 8.4 |
| C_0021 | Creatine | Creatine | 586 | 23 | 11 | 15 | 34 | 2.0 | 4.1 | 61 | 3.3 | 3.9 |
| C_0022 | Ile | Ile | 791 | 11 | 40 | 56 | 218 | 225 | 290 | 315 | 265 | 602 |
| C_0023 | Leu | Leu | 857 | 14 | 42 | 59 | 207 | 213 | 262 | 293 | 248 | 574 |
| C_0024 | Asn | Asn | 236 | 5.4 | 36 | 48 | 185 | 104 | 189 | 173 | 112 | 525 |
| C_0025 | Ornithine | Ornithine | 389 | 1.4 | 1.3 | N.D. | 22 | 15 | 14 | 29 | 15 | 40 |
| C_0026 | Asp | Asp | 424 | 783 | 978 | 436 | 1,977 | 1,975 | 2,289 | 358 | 518 | 1,437 |
| C_0027 | Homocysteine | Homocysteine | 778 | N.D. | N.D. | N.D. | N.D. | N.D. | N.D. | N.D. | N.D. | N.D. |
| C_0028 | Adenine | Adenine | 190 | 1.0 | 1.0 | N.D. | N.D. | N.D. | N.D. | 0.7 | N.D. | N.D. |
| C_0029 | Hypoxanthine | Hypoxanthine | 790 | N.D. | N.D. | N.D. | N.D. | N.D. | N.D. | 2.6 | N.D. | 4.7 |
| C_0030 | Spermidine | Spermidine | 1102 | N.D. | N.D. | N.D. | N.D. | N.D. | 4.0 | 2.4 | 1.5 | 2.2 |
| C_0031 | Gln | Gln | 738 | 52 | 63 | 67 | 1,275 | 1,203 | 1,906 | 576 | 792 | 1,892 |
| C_0032 | Lys | Lys | 866 | 55 | 62 | 45 | 175 | 181 | 171 | 145 | 127 | 376 |
| C_0033 | Glu | Glu | 611 | 122 | 713 | 617 | 3,875 | 3,388 | 5,976 | 3,273 | 2,947 | 6,057 |
| C_0034 | Met | Met | 876 | 5.0 | 14 | 18 | 39 | 48 | 55 | 53 | 38 | 94 |
| C_0035 | Guanine | Guanine | 764 | N.D. | N.D. | N.D. | N.D. | N.D. | N.D. | N.D. | N.D. | N.D. |
| C_0036 | His | His | 773 | 11 | 18 | 19 | 66 | 63 | 78 | 66 | 61 | 137 |
| C_0037 | Carnitine | Carnitine | 85 | 17 | 11 | 17 | N.D. | N.D. | N.D. | N.D. | N.D. | N.D. |
| C_0038 | Phe | Phe | 994 | 10 | 22 | 26 | 82 | 97 | 107 | 135 | 122 | 280 |
| C_0039 | Arg | Arg | 6322 | 31 | 31 | 34 | 143 | 113 | 88 | 92 | 49 | 161 |
| C_0040 | Citrulline | Citrulline | 9750 | 4.0 | 3.0 | 2.8 | 4.9 | N.D. | N.D. | 2.3 | N.D. | N.D. |
| C_0041 | Tyr | Tyr | 1153 | 12 | 21 | 26 | 73 | 80 | 88 | 129 | 112 | 252 |
| C_0042 | *S*-Adenosylhomocysteine | SAH | 439155 | N.D. | N.D. | N.D. | N.D. | N.D. | N.D. | 1.3 | N.D. | N.D. |
| C_0043 | Spermine | Spermine | 1103 | N.D. | N.D. | N.D. | N.D. | N.D. | 13 | 9.3 | 5.5 | 6.2 |
| C_0044 | Trp | Trp | 1148 | N.D. | 5.8 | 5.7 | 19 | 19 | 18 | 29 | 24 | 57 |
| C_0045 | Cystathionine | Cystathionine | 834 | N.D. | N.D. | N.D. | N.D. | N.D. | N.D. | N.D. | N.D. | 3.4 |
| C_0046 | Adenosine | Adenosine | 60961 | N.D. | N.D. | N.D. | N.D. | N.D. | 3.0 | 1.5 | N.D. | N.D. |
| C_0047 | Inosine | Inosine | 6021 | N.D. | N.D. | N.D. | N.D. | N.D. | N.D. | N.D. | N.D. | N.D. |
| C_0048 | Guanosine | Guanosine | 6802 | N.D. | N.D. | N.D. | N.D. | N.D. | N.D. | N.D. | N.D. | N.D. |
| C_0049 | Argininosuccinic acid | ArgSuccinate | 16950 | N.D. | N.D. | N.D. | N.D. | N.D. | N.D. | 5.7 | 3.3 | 8.5 |
| C_0050 | Glutathione (GSSG) | GSSG | 65359 | 115 | 132 | 80 | 189 | 377 | 151 | 424 | 774 | 694 |
| C_0051 | Glutathione (GSH) | GSH | 124886 | 101 | 108 | 122 | 216 | 83 | 1,118 | 968 | 970 | 2,237 |
| C_0052 | *S*-Adenosylmethionine | SAM | 34755 | 8.2 | 4.9 | 3.9 | 12 | N.D. | 8.9 | 9.8 | 9.3 | 27 |
| - | Adenylate Energy Charge | No Label | | 0.7 | 0.9 | 0.9 | 1.0 | 1.0 | 0.9 | 0.8 | 0.9 | 0.8 |
| - | Total Adenylate | No Label | | 251 | 328 | 307 | 1,694 | 1,173 | 2,277 | 2,285 | 1,510 | 2,271 |
| - | Guanylate Energy Charge | No Label | | 0.7 | 0.9 | 0.9 | 1.0 | 0.9 | 0.9 | 0.8 | 0.9 | 0.9 |
| - | Total Guanylate | No Label | | 62 | 74 | 62 | 298 | 201 | 419 | 580 | 352 | 504 |
| - | GSH/GSSG | No Label | | 0.9 | 0.8 | 1.5 | 1.1 | 0.2 | 7.4 | 2.3 | 1.3 | 3.2 |
| - | Total Glutathione | No Label | | 332 | 372 | 282 | 593 | 838 | 1,420 | 1,815 | 2,519 | 3,624 |
| - | NADPH/NADP+ | No Label | | 3.3 | 4.5 | 4.4 | 3.2 | 3.4 | 1.9 | 2.0 | 1.8 | 2.0 |
| - | NADH/NAD+ | No Label | | 1.2 | 0.7 | 0.6 | 0.09 | 0.14 | 0.06 | 0.09 | 0.09 | 0.10 |
| | NAD+/NADH | | | 0.9 | 1.3 | 1.7 | 11.3 | 7.0 | 15.7 | 11.1 | 11.6 | 10.2 |
| - | Lactate/Pyruvate | No Label | | 3.1 | 5.1 | 6.7 | 24 | 22 | 24 | 112 | 98 | 61 |
| - | Glycerol 3-phosphate/DHAP | No Label | | 1.0 | 1.4 | 13 | N.A. | N.A. | 122 | 217 | 67 | 39 |
| - | Total Amino Acids | No Label | | 1,409 | 2,691 | 2,160 | 11,878 | 10,337 | 16,132 | 10,614 | 8,678 | 19,946 |
| - | Total Essential Amino Acids | No Label | | 162 | 324 | 364 | 1,393 | 1,466 | 1,964 | 1,814 | 1,684 | 3,866 |
| - | Total Non-essential Amino Acids | No Label | | 1,247 | 2,368 | 1,796 | 10,486 | 8,871 | 14,168 | 8,800 | 6,994 | 16,080 |
| - | Total Glucogenic Amino Acids | No Label | | 1,340 | 2,587 | 2,056 | 11,496 | 9,943 | 15,698 | 10,177 | 8,303 | 18,997 |
| - | Total Ketogenic Amino Acids | No Label | | 157 | 275 | 301 | 1,141 | 1,206 | 1,568 | 1,510 | 1,411 | 3,252 |
| - | Total BCAA | No Label | | 25 | 120 | 167 | 643 | 667 | 903 | 921 | 798 | 1,812 |
| - | Total Aromatic Amino Acids | No Label | | 22 | 48 | 58 | 173 | 196 | 213 | 293 | 259 | 590 |
| - | Fischer's Ratio | No Label | | 1.2 | 2.5 | 2.9 | 3.7 | 3.4 | 4.2 | 3.1 | 3.1 | 3.1 |
| - | Total Glu-related Amino Acids | No Label | | 258 | 954 | 875 | 5,716 | 4,911 | 8,891 | 5,790 | 4,806 | 10,151 |
| - | Total Pyr-related Amino Acids | No Label | | 256 | 486 | 519 | 2,989 | 2,274 | 3,438 | 2,911 | 2,045 | 5,019 |
| - | Total Acetyl CoA-related Amino Acids | No Label | | 80 | 150 | 166 | 618 | 639 | 741 | 781 | 664 | 1,609 |
| - | Total Fumarate-related Amino Acids | No Label | | 22 | 42 | 52 | 154 | 176 | 196 | 263 | 235 | 532 |
| - | Total Succinyl CoA-related Amino Acids | No Label | | 16 | 91 | 126 | 475 | 503 | 696 | 681 | 588 | 1,333 |
| - | Total Oxaloacetate-related Amino Acids | No Label | | 788 | 1,013 | 483 | 2,161 | 2,079 | 2,478 | 532 | 630 | 1,962 |
| - | Malate/Asp | No Label | | 0.07 | 0.03 | 0.05 | N.A. | N.A. | 0.02 | 0.9 | 0.5 | 0.2 |
| - | Citrulline/Ornithine | No Label | | 2.9 | 2.4 | N.A. | 0.2 | N.A. | N.A. | 0.08 | N.A. | N.A. |
| - | Glu/2-Oxoglutarate | No Label | | N.A. | N.A. | N.A. | N.A. | N.A. | N.A. | 52 | 132 | 205 |
| - | G6P/R5P | No Label | | 1.2 | 2.2 | 2.1 | 4.6 | 3.0 | 0.7 | 2.2 | 7.9 | 2.5 |
| - | SAM/SAH | No Label | | N.A. | N.A. | N.A. | N.A. | N.A. | N.A. | 7.4 | N.A. | N.A. |
| - | Putrescine/Spermidine | No Label | | N.A. | N.A. | N.A. | N.A. | N.A. | N.A. | N.A. | N.A. | N.A. |

ID consists of analysis mode and number. 'C' and 'A' showed cation and anion modes, respectively.
N.D. (Not Detected): The target peak or metabolite was below detection limits.
N.A. (Not Available): The calculation was impossible because of insufficience of the data.¶
The data are sorted by ID in ascending order.
